# Forest bird decline and community change over 19 years in long-isolated South Asian tropical rainforest fragments

**DOI:** 10.1101/2022.10.22.513365

**Authors:** Akshay Surendra, T. R. Shankar Raman

**Affiliations:** Nature Conservation Foundation, 1311, 12^th^ A Main, Vijayanagar 1^st^ Stage, Mysore 570017, India; School of the Environment, Yale University, New Haven, CT – 06511, USA; New York Botanical Garden, 2900 Southern Blvd, Bronx, NY 10458

**Keywords:** forest fragmentation, habitat structure, patch size, Western Ghats, tropical bird conservation

## Abstract

Recent evidence of forest bird declines worldwide is attributed to climate change and its interactive effects with recent land-use changes such as forest loss and fragmentation, and avian life-history traits. In Asian tropical forests, such effects are poorly understood as long-term data are lacking from fragments that are long-isolated rather than recently fragmented. Here, we use data from ~2000 point-counts from bird surveys carried out between 2000 – 2005 and 2019 in 19 long-isolated (~80 y) South Asian tropical rainforest fragments to examine changes in bird species richness, density, and composition in relation to fragment area (0.7 – 4310 ha), habitat structure, and time. Over the 19 y timespan, despite stable fragment areas, we uncovered a 29% decline in rainforest bird density and 7% decline in individual-rarefied species richness of rainforest birds, while density and richness of open-country birds remained stable. With increasing fragment area, rainforest bird species richness (jackknife estimate) increased, while open country bird richness (individual-rarefied) and density decreased. Larger fragments housed more compositionally stable bird communities, while poorer habitat was associated with lower diversity of rainforest birds but higher diversity, density, and compositional variation of open-country birds. Threshold analysis however indicated relatively small area thresholds (~20 ha) for rainforest bird species abundance. Besides identifying alarming declines in rainforest birds, the study confirms some but not all predictions for bird diversity in long-isolated forest fragments with stable forest-matrix boundaries, indicating that small fragments and habitat quality also matter.

## Introduction

Pervasive declines of bird diversity and abundance have been reported from tropical forests around the world (Sherry 2021). Such decreases at local scales have been attributed to lag-effects from proximate land-use change, such as extinction debt from forest loss and fragmentation (Tilman *et al.* 1994). However, the decline in bird populations at the scale of continents (Bowler *et al.* 2019, Rosenberg *et al.* 2019), including within undisturbed habitats like in Amazonian rainforest (Stouffer *et al.* 2021) point to the putative role of global and regional stressors linked to climate change. The overall change in bird populations is likely to be the result of recent and historic anthropogenic disturbances at both local (Laurance 2008) as well as global or regional changes in climate (Sherry 2021, Lees *et al.* 2022). Studies that fail to detect population declines nevertheless reveal some form of compensatory buffering: generalist birds make up for declining specialists (Bowler *et al.* 2019), taxonomic redundancy buffers against the erosion of functional diversity (Oliveira & dos Anjos 2022), and even within a species, birds adapt (Jirinec *et al.* 2021) or acclimatize (Martin & Mouton 2020) to increasing stress from changing land-use and climate. Current understanding of such synergistic climate and land-use effects on bird communities (Srinivasan & Wilcove 2021) is constrained by sparse long-term data (Robinson & Curtis 2020).

Consider forest fragmentation after forest loss – a ubiquitous and persistent threat to forests and their biodiversity across the planet (Fahrig 2017, Taubert *et al.* 2018). Over the last few decades, ‘snapshot studies’ across ecosystems have revealed some well-understood and generalisable patterns in bird community responses to fragmentation (Laurance 2008). Studies indicate that large, well-connected habitat patches within a low-contrast, high habitat-quality matrix are better for bird diversity than small, more-isolated habitat patches within a high-contrast, low habitat-quality matrix (Turner 1996, Prevedello & Vieira 2010). Also, edge effects have a well-understood (largely) adverse impact on birds (Harper *et al.* 2005), although their responses to forest edges and fragmentation may be modulated by species traits (Henle *et al.* 2004, Bregman *et al.* 2014). Our understanding of temporal changes in bird communities across fragmented landscapes comes from fewer studies (Debinski & Holt 2000). Long-term fragmentation research reveals that birds initially compress into habitat patches from recently deforested landscape matrices (Stouffer 2020), following which populations decline rapidly towards stable abundance (Brotons *et al.* 2003) while richness declines exponentially (Stouffer *et al.* 2009). However, these long-term studies – despite their importance – only characterise transient and not equilibrium dynamics within fragmented forests (Luther *et al.* 2020).

Theory predicts a dynamic equilibrium over time (Brooks *et al.* 1999) of constant species richness in forest fragments where recolonisation (species credit (Lira *et al.* 2012, Latta *et al.* 2017)) catches up with declining richness (extinction debt (Tilman *et al.* 1994)). Thus, long-isolated forest patches with stable habitat-matrix boundaries are likely to be maintained as a stable meta-population (Rolstad 1991) with relatively constant richness and abundance (but not composition) of birds. On the other hand, if species richness, abundance, or both continue to decline in such long-isolated fragments over time, global or regional drivers are likely at play – singly or in tandem with land-use change (Sigel *et al.* 2010). While a handful of studies reveal species declines long after fragmentation (e.g., (Sigel *et al.* 2010, Curtis *et al.* 2021, Pollock *et al.* 2022)), generalisations or region-specific trends are difficult to identify because of the near absence of long-term studies of birds from tropical forest fragments outside the Neotropics (Fardila *et al.* 2017).

As forests around the world continue to become more fragmented (Haddad *et al.* 2015, Taubert *et al.* 2018), the ability of forest fragments to house stable populations of forest-specialist bird species across varying fragment sizes – in other words, the conservation value of remnant forest patches (Turner & Corlett 1996) – remains crucial to establish.

Here, we report patterns in density, richness, and composition of bird communities across 19 years within a long-isolated fragmented rainforest landscape in southern India (Raman & Mudappa 2003, Raman 2006, Anand *et al.* 2010). We disaggregate the bird community into habitat-specialist rainforest (henceforth, RF) birds or matrix-derived open-country (henceforth, OC) birds in order to identify potentially contrasting effects from these two broad guilds (Raman 2006). We also subset birds with restricted home-ranges (henceforth, RR) that may be particularly vulnerable due to poorly-defined (Ramesh *et al.* 2017) or potentially, poorly-protected (Cazalis *et al.* 2021) species ranges. We separately examine all three subsets, as well as the entire bird community taken together (all birds).

We expected that stable bird density and richness would be consistent with a dynamic equilibrium long after fragmentation, with no discernible deleterious effects of regional climate stressors. On the other hand, declines in richness or density would indicate a putative link with increasing regional stressors. Monsoon rains in the southern Western Ghats have steadily declined in the past century (Varikoden *et al.* 2019) while post-monsoon cyclonic events in the Bay of Bengal have increased in frequency and intensity (Balaguru *et al.* 2014). Unless birds and the vegetation they depend on rapidly acclimate (Martin & Mouton 2020) or adapt (Van Buskirk *et al.* 2010) to such shifts in climate, both the density and diversity of bird assemblages may decline (Ramesh *et al.* 2022). (Ramesh *et al.* 2022)Species with restricted ranges (Robin *et al.* 2015) and narrower extant habitat and climatic envelops are likely to be especially sensitive to such climate change (Şekercioğlu *et al.* 2012). Any change in bird communities through time would be modulated by habitat: fragments in our study system have idiosyncrasies specific to each fragment, including higher than average secondary disturbance (for instance, invasive understory shrubs (Joshi *et al.* 2009), preferential extraction of pole-sized trees (Mudappa & Raman 2007), or active or passive restoration (Osuri *et al.* 2019) – all of which can alter understory structure over two decades.

We use six annual point count surveys across 19 fragments in 2000-05 and 2019 to test patterns in diversity, density, and composition through time, while controlling for the effect of fragment size and changing habitat quality. We identify species-specific abundance thresholds in relation to fragment area to pinpoint area-sensitive taxa – bird species whose abundance decreases or increases substantially beyond the thresholds – and their implications for patch sizes relevant for bird conservation.

## Methods

### Study area

The study area in the Anamalai Hills, a significant conservation area in southern India's Western Ghats Biodiversity Hotspot (Kumar *et al.* 2004), included forest fragments in the Valparai Plateau (220 km^2^, 10.30° N, 76.95° E) and intact forests in the adjoining Anamalai Tiger Reserve (958 km^2^, Figure 1). Annual rainfall averages about 2400 mm, over two-thirds of which falls during the southwest monsoon (June to September). Within the elevation range of 700 m – 1500 m ASL, the natural vegetation, confined to about 45 forest fragments within the plateau and adjoining tiger reserve, is characterised as mid-elevation tropical wet evergreen rainforests of the *Cullenia exarillata* – *Mesua ferrea* – *Palaquium ellipticum* type (Pascal *et al.* 2004). The matrix around the fragments consists of tea and coffee plantations established between 1890 and 1920 during the British colonial period and currently owned by large plantation companies. These forest fragments have persisted in the plantation land-use matrix with limited disturbance and relatively steady matrix-habitat boundaries for at least seven decades (Mudappa & Raman 2007). The Valparai plantation landscape also supports a human population of around 70,000 people, largely plantation workers and indigenous people who depend on forests in the Anamalai Tiger Reserve for their livelihood. Further details of the study area, including more landscape information and an outline of its ecological history, can be found elsewhere (Mudappa & Raman 2007).

**Figure 1:**
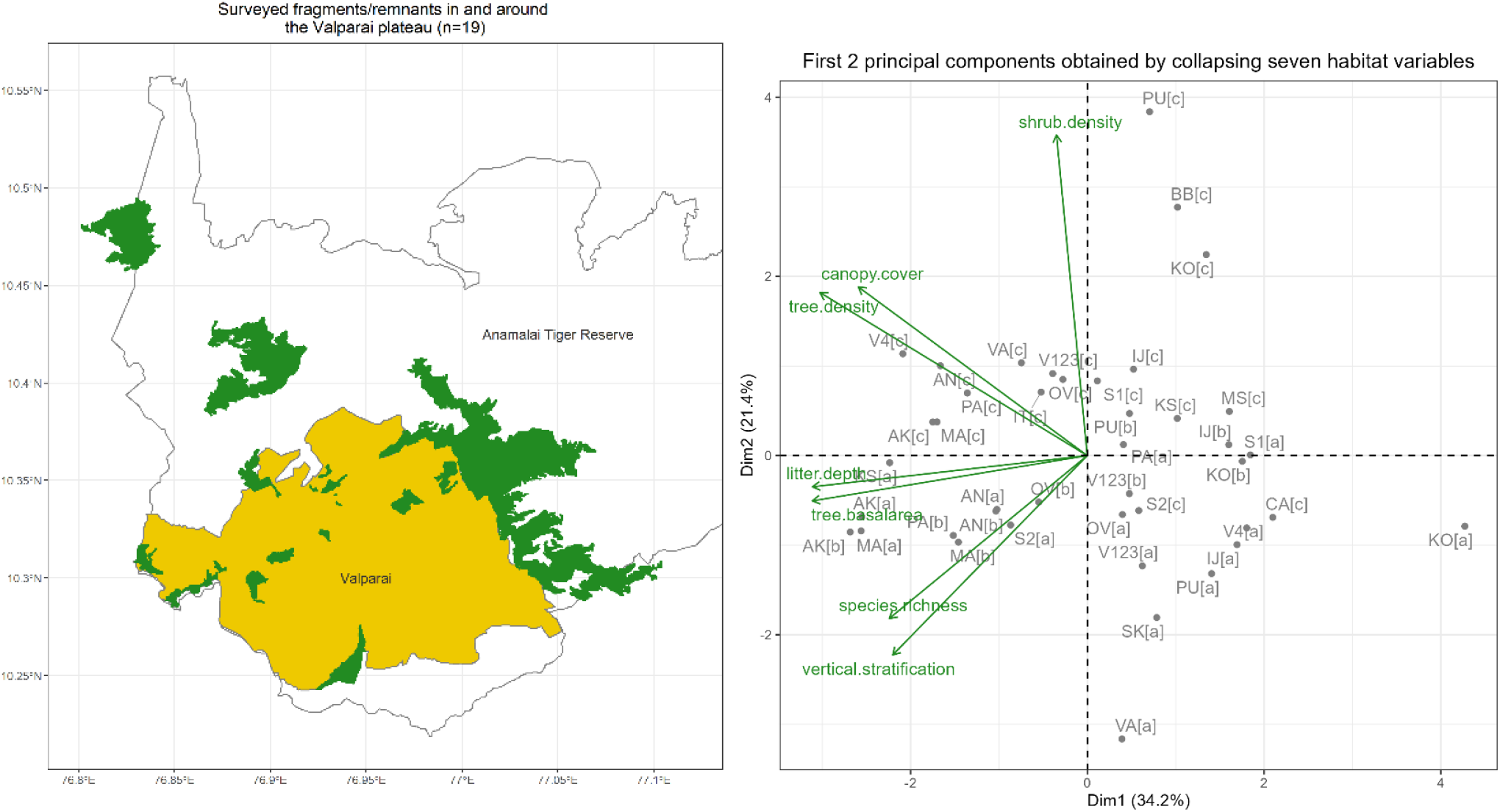
a) Forest fragments (dark green) within and around the Valparai Plateau plantation landscape (yellow) in the Anamalai Hills, Western Ghats, India. b) Bi-plot of the first two dimensions from a Principal Components Analysis (PCA) of habitat variables (see Table S2 for eigenvalues and axes loadings). Points indicate vegetation characteristics in each fragment (codes as in Table 1), with [a], [b], and [c] indicating sampling years 2000, 2005, and 2019, respectively.

We sampled bird communities and vegetation in 19 rainforest fragments ranging in size from 0.67 ha to 4310 ha (Figure 1). These included 13 fragments (0.67 to 113 ha) embedded within tea and coffee plantations on the Valparai Plateau and six fragments (175 to 4310 ha) in the Anamalai Tiger Reserve that also abutted montane grasslands or deciduous forests (Osuri *et al.* 2020). The spatially heterogeneous landscape also has riparian forests, small abandoned plantations, and a few ecologically restored fragments, which enhance landscape-level forest connectivity (Wordley *et al.* 2015). Due to this and because all forest fragments were <0.5 km away from other forest patches and <3 km from continuous forests, we did not consider fragment isolation as a potential driver of bird community structure in our study.

**Table 1:** Sampling effort, fragment characteristics and mean ± SE of estimated bird density (birds per hectare) and individual rarefied species richness (species per hectare, rarefied to 46 individuals) across 19 sampled fragments

| Fragment | Fragment area (ha) | Total Point Counts | Years sampled /total years | Total number of bird species | Bird density (ha<sup>-1</sup>) | Rarefied species richness (N<sub>min</sub> = 46) |
| --- | --- | --- | --- | --- | --- | --- |
| Akkamalai Iyerpadi (AK) | 4309.98 | 433 | 6/6 | 70 | 25.2 ± 4 | 10.5 ± 1.7 |
| Varagaliar (VA) | 1411.09 | 82 | 2/6 | 61 | 15.1 ± 4.5 | 11.7 ± 3.2 |
| Karian Shola KS) | 854.27 | 99 | 2/6 | 64 | 14.4 ± 4.1 | 11.5 ± 3.1 |
| Andiparai (AN) | 327.00 | 181 | 4/6 | 61 | 22.6 ± 5.1 | 10.2 ± 2 |
| Sankarankudi (SK) | 252.49 | 25 | 1/6 | 45 | 21.6 ± 7.1 | 11.4 ± 5.1 |
| Manamboly (MA) | 174.95 | 212 | 5/6 | 84 | 24.4 ± 4.3 | 11.8 ± 2.1 |
| Puduthottam (PU) | 113.08 | 239 | 6/6 | 77 | 21.2 ± 2.5 | 11 ± 1.8 |
| Iyerpadi Top (IT) | 103.91 | 44 | 1/6 | 49 | 22.4 ± 5.8 | 9.9 ± 4.4 |
| Candura (CA) | 103.30 | 58 | 1/6 | 59 | 29.1 ± 9.1 | 11.1 ± 5 |
| Murugalli Sholayar (MS) | 102.43 | 50 | 1/6 | 67 | 31.4 ± 9 | 11.7 ± 5.3 |
| Pannimade (PA) | 92.13 | 128 | 3/6 | 70 | 29.3 ± 6.4 | 10.9 ± 2.6 |
| Korangumudi (KO) | 73.32 | 203 | 5/6 | 94 | 24.4 ± 3.6 | 11.7 ± 2.1 |
| Old Valparai (OV) | 41.31 | 172 | 5/6 | 81 | 27.2 ± 3.8 | 11.4 ± 2 |
| Injipara (IJ) | 18.98 | 145 | 5/6 | 80 | 26.6 ± 3.3 | 10.8 ± 1.9 |
| Murugalli Black Bridge (BB) | 15.79 | 20 | 1/6 | 41 | 34.2 ± 11 | 11 ± 5 |
| Varattuparai123 (V123) | 12.52 | 74 | 3/6 | 74 | 23 ± 4.5 | 11 ± 2.6 |
| Varattuparai4 (V4) | 3.85 | 21 | 2/6 | 42 | 24.1 ± 6.5 | 9.8 ± 2.9 |
| Selaliparai1 (S1) | 2.22 | 23 | 2/6 | 48 | 31.6 ± 7.3 | 10.9 ± 3.2 |
| Selaliparai2 (S2) | 0.67 | 14 | 2/6 | 44 | 30.7 ± 7.8 | 12.3 ± 3.6 |

### Bird Surveys

We sampled the 19 forest fragments between 2000 and 2005 and in 2019. TRSR sampled between 2000 to 2004, Hari Sridhar (HS) in 2005 and AS in 2019. HS and AS conducted initial bird surveys jointly with TRSR to ensure that all observers used similar methods and had similar bird identification skills. Each sampling season in the designated year extended from the previous December to early May, when breeding residents and long-distance migrants occur in the area (Raman 2006). We conducted 2223 point count surveys, averaging 371 point counts per season and 39 point counts per fragment per season. Overall, we surveyed 14 fragments in at least two seasons, nine fragments in at least three, and seven were surveyed across four or more seasons (see Table 1 for sampling effort and fragment characteristics).

These variable-radius point count surveys were carried out during the first three hours after sunrise – when bird activity is highest – following standard protocols (Raman 2003, Buckland *et al.* 2015). We identified and counted all birds seen or heard within replicate 5-min duration surveys; all detections were binned into five radial distance classes (0 – 5, 5 – 10, 10 – 20, 20 – 30, 30 – 50: in metres) that were estimated visually (if seen) or aurally (if heard). We spaced successive point count locations by at least 100 m when sampled on the same day (Raman 2006). We revisited sites after a gap of 3 – 4 weeks. We covered different parts of the forest fragment on each visit as far as possible to capture within-season and within-fragment heterogeneity in bird communities while minimising spatial and temporal autocorrelation These data were first uploaded to the eBird India portal (http://ebird.org/india) to pass the citizen science review process.

### Vegetation sampling

We measured vegetation characteristics in 2000, 2005, and 2019 using standard field methods for relevant habitat variables used in earlier studies (Raman *et al.* 1998, Raman 2001b, 2006, Sridhar & Sankar 2008). To estimate the density, basal area, and rarefied species richness of trees (>30 cm girth at breast height), we used data from 15–30 point-centred quarter (PCQ) samples spaced about 100 m apart in each fragment (Mitchell 2015), noting tree girth, radial distance from the point, and species identity (identity not recorded in 2005). We also measured four habitat variables—canopy cover, vertical stratification, shrub density, and leaf-litter depth—at each PCQ location. We measured canopy cover using a spherical densiometer and vertical stratification by summing the number of presences (1) or absences (0) of foliage in eight vertical bands (0 – 1, 1 – 2, 2 – 4, 4 – 8, 8 – 16, 16 – 24, 24 – 32: in meters) directly overhead. We measured leaf litter depth to nearest 0.5 cm using a ruler. The seven habitat variables served as proxies of rainforest habitat quality, with larger values tending to be associated with more species-rich and abundant rainforest bird communities (Raman *et al.* 1998, Raman 2001b, a, 2006).

### Analysis

We tested our hypotheses on four different sets of birds’ point count data: all birds (All), rainforest birds (RF), open-country birds (OC) and range-restricted species (RR). RF and OC are mutually exclusive subsets of the larger dataset of all birds; RR species are a subset of rainforest birds that represent birds endemic to the Western Ghats or extending into adjoining hill ranges of peninsular India (Raman 2006). Bird handbooks and field guides were used to assign species to these habitat groupings in the region (Ali *et al.* 1983). Two pairs of sister taxa were considered single species as taxonomic splits had not occurred in the earlier surveys (e.g., Green Warbler *Phylloscopus nitidus* and Greenish Warbler *P. trochiloides*, and Chestnut-tailed Starling *Sturnus malabarica* and Malabar Starling *S. blythii*).

We used the R programming language (version 4.0.2 (R Core Team 2013)) for our analysis. In particular, we used package tidyverse (version 1.3.0 (Wickham 2017)) for data management, package factoextra (version 1.0.7 (Kassambara & Mundt 2017)) to visualise the PCA bi-plot, package Distance (version 1.0.0 (Miller *et al.* 2016)) to estimate bird density, package vegan (version 2.5-6 (Oksanen *et al.* 2007)) to compute species richness and compositional dissimilarity, lmerTest (version 3.1-3 (Kuznetsova *et al.* 2017)) to run our models and sjPlot (version 2.8.5 (Lüdecke 2018b)) to make forest plots and ggeffects (version 1.1.3 (Lüdecke 2018a)) to calculate effect sizes and make effect plots.

Bird species richness was estimated through rarefaction (for a standardised number of individuals to account for density-differences across years): all birds, rainforest birds, open-country birds, and range-restricted birds were rarefied by sampling with replacement 46, 23, 3, and 8 individuals, respectively, and estimating unique species from this bootstrapping procedure (Gotelli & Colwell 2001). We estimated asymptotic bird species richness using a pooled first-order jackknife estimate (Gotelli & Chao 2013).

Bird density was estimated from the variable-radius point counts data using distance sampling for all seasons except 2005 when detection detections were not evaluated in the field (Raman 2003, Buckland *et al.* 2015). We used estimated and not exact distances because, typical of the tropics (Robinson *et al.* 2018), 88% of all detections were heard and not seen. Density was estimated for each fragment, taking each point count as a spatial replicate, accounting for two species covariates—‘loudness’ or ease of aural detection and ‘sightability’ or ease of visual detection (further details in Supplementary Table S3).

Compositional stability was measured by computing pairwise dissimilarity of each fragment with itself over time (Magurran *et al.* 2010). We used the Bray-Curtis dissimilarity index over a Hellinger-transformed species-site matrix with species as columns, each fragment-year combination as rows, and populating this matrix with total detections: we excluded fragments that were sampled only once in the 19 years.

Time, fragment area, and season-specific habitat quality were used as putative drivers of bird community structure. In models with species richness and bird density as the response, we used sampling year as the time variable; in models with compositional dissimilarity as the response, we used the time difference (in years) between the two sampling seasons. Forest fragments were mapped using Google Earth and a handheld GPS to obtain boundaries and estimate fragment size.

We summarised variation in seven habitat quality variables—tree density, tree basal area, tree species richness, vertical stratification, canopy cover, leaf-litter depth, and shrub density—across 42 fragment – season combinations using Principal Component Analysis. As vegetation was not sampled between 2001 and 2004, we assigned habitat predictor estimates from the nearest season (either 2000 or 2005), assuming low turnover in trees and habitat structure across such short time intervals. As leaf-litter depth and tree species identity were not recorded in 2005, we used corresponding estimates from 2000. We retained the first three principal component axes (PC1 to PC3) that together explained 69.7% of the total variance (Figure 1b, Table S2).

The effects of time, fragment size, and habitat quality on bird community structure were assessed using Generalised Linear Mixed Models (GLMMs) (Bates *et al.* 2015). We separately regressed species richness, bird density, and compositional dissimilarity for each bird guild (all, rainforest, open-country, and range-restricted) against time (or time-span), fragment size, and the first three PC axes representing habitat quality. Since several fragments were sampled more than once, we used fragment identity as a random intercept to account for repeat surveys. We scaled and centred all predictors before using them in the model, and all sets of predictors were included after checking for multi-collinearity. We used identity links for all three responses and achieved robust model fits for all except for open-country subset’s density model. We used a simple linear model instead, using leave-one-out regression with random effects to verify that direction and significance of all predictors were consistent across all the models with or without Fragment as a random effect.

Finally, we examined fragment size thresholds for species abundance responses using Threshold Indicator Taxon ANalysis (or TITAN) (Baker & King 2010) implemented through an eponymous package in R (Baker 2019). TITAN uses indicator value scores (Dufrene & Legendre 1997) of species, creating binary partitions along environmental (in this case, fragment size) gradients to detect congruent changes in species-specific abundance and occurrence frequency and identify change points as potential ecological thresholds. To include information from 2005 (when bird density was not estimated), we replaced abundances for all detections (including 2005) with unity: this represents a reasonably conservative estimate since 77.3% of all bird detections across the other sampling seasons were singletons. This measure of species abundance is analogous to occurrence frequency (or frequency-of-checklists metric used in the citizen science portal eBird) (Sullivan *et al.* 2009, SoIB 2020)). We subset species that occurred at least thrice in each of the 57 fragment-year combinations to satisfy TITAN’s minimum count requirement (Baker 2019). Preliminary analysis revealed that most species’ thresholds lay below 300 hectares: those with more significant fragment thresholds were high-elevation specialists (for example, White-bellied Sholakili), and only two of our larger fragments (Akkamalai-Iyerpadi and Andiparai) comprised of habitat suitable for such species, precluding the ability to disentangle the effect of fragment area from elevation. Thus, we restricted TITAN to 15 small and medium sized fragments (of area < 300 ha).

## Results

We recorded 23,176 detections in our sample of 2,223 point-counts across 19 years and 19 fragments (Table 1, Figure 1a). We identified 99.15% of these bird detections to the species level, and this included a total of 131 species: 92 RF (including 22 RR), and 39 OC species (Table S1 lists species, their corresponding habitat guilds, and detections over the 19-year period).

Habitat structure differed among fragments (Figure S1) as revealed by principal components analysis (PCA) of habitat variables (Figure 1b). The PC1 axis explained 34.2% of the total variance in habitat variables. Higher PC1 scores were mainly associated with lower tree density, canopy cover, and leaf litter depth (Figure 1b, Table S2). The PC2 axis explained 21.4% of the total variance. Higher PC2 scores were mainly associated with greater shrub density in the understory and reduced vertical stratification (Figure 1b, Table S2). The PC3 axis explained 14.0% of the total variance. Higher PC3 scores were associated with greater tree basal area and lower tree species richness (Table S2). In general, larger and less disturbed fragments had lower PC1 scores (Figure 1b).

### Bird community structure over time

Rarefied species richness of rainforest and restricted-range birds declined by 7% and 14% over 19 years (*P* < 0.05) while open-country birds declined marginally (*P* = 0.07) by 12% over the same period (Figure 2, Table 2). Jackknife species richness remained stable for all except range-restricted birds, which showed a marginal increase (19% over 19 years, *P* = 0.09). We assessed the effects of time on bird abundance using density estimates from distance sampling (Table S3). The density of rainforest birds declined by 29% over 19 years (*P* < 0.05) but the density of range-restricted and open-country birds remained stable (*P* = 0.54 and *P* = 0.88, Table 2). Dissimilarity in species composition predictably increased over longer intervals for all three habitat guilds (*P* < 0.05, Table 2). The overall pattern for all-birds reflected the net effect of all guilds: the density of all birds taken together declined over time (*P* < 0.05), compositional dissimilarity increased with time (*P* < 0.05), rarefied species richness decreased marginally (*P* = 0.07), and jackknife richness increased marginally (*P* = 0.07).

**Table 2:**
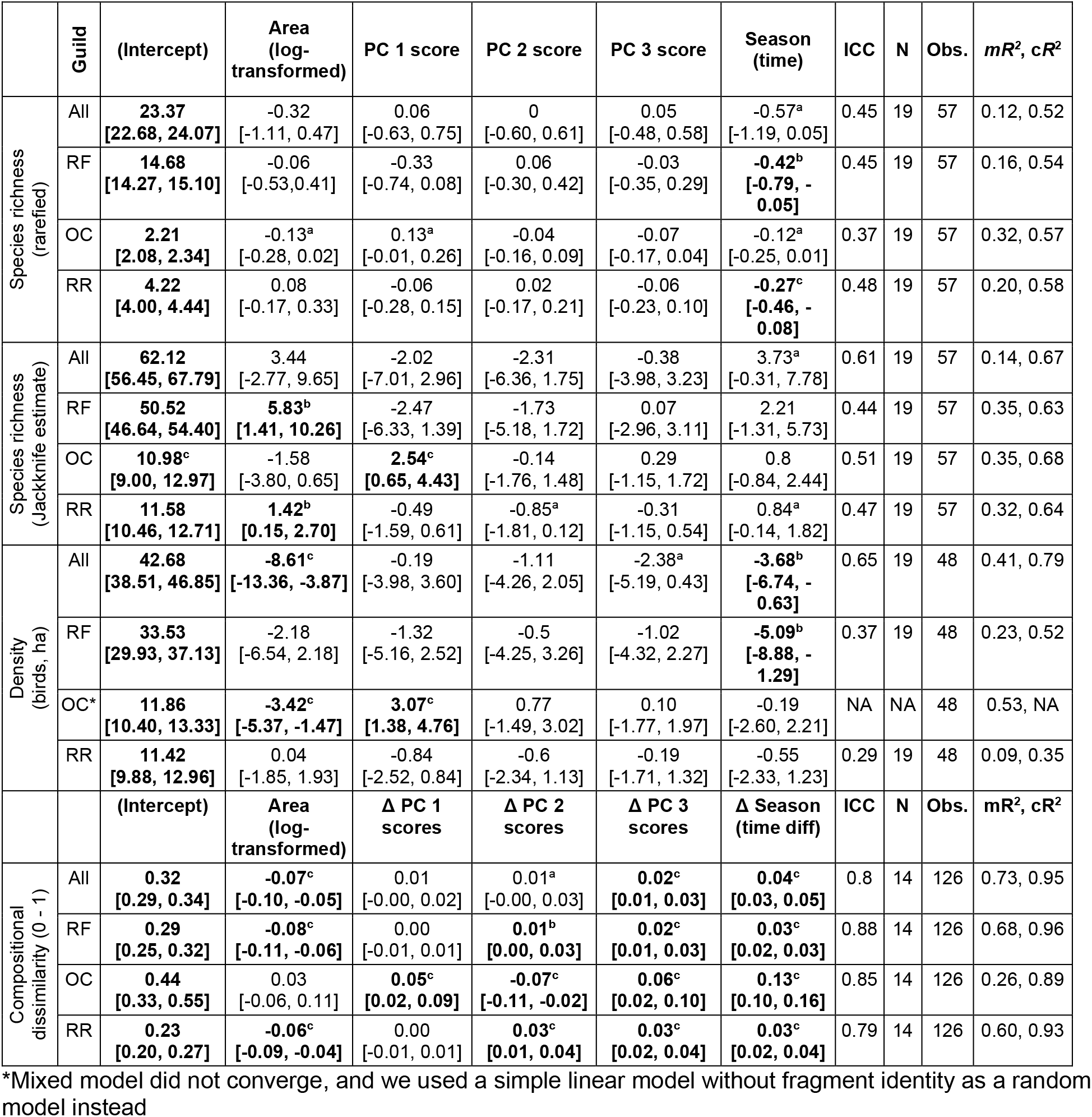
Linear mixed model regressions of bird species richness (rarefaction and jackknife estimates), bird density (distance sampling estimate), and compositional dissimilarity of bird communities (Bray-Curtis dissimilarity) of birds against time, fragment area, and habitat quality as indexed by Principal Component (PC) scores; fragment identity is the random effect and the intraclass correlation coefficient (ICC) is the variance fraction explained by fragment identity. Tabled values indicate regression coefficients (95% confidence interval), sample size of fixed effects (N = 19 or 14 fragments) and random levels (Observations = 57, 48 or 126), and marginal (m*R*^2^) and conditional (c*R*^2^) R-squared. [All = entire bird community; RF = rainforest birds; RR = restricted-range birds; OC = open-country birds]. Statistically significant coefficients are indicated in bold: ^a^ – *P* < 0.10, ^b^ – *P* < 0.05, ^c^ – *P* < 0.01.

| | Guild | (Intercept) | Area (log-transformed) | PC 1 score | PC 2 score | PC 3 score | Season (time) | ICC | N | Obs. | $mR^2$, $cR^2$ |
| --- | --- | --- | --- | --- | --- | --- | --- | --- | --- | --- | --- |
| Species richness (rarefied) | All | **23.37**<br>[22.68, 24.07] | -0.32<br>[-1.11, 0.47] | 0.06<br>[-0.63, 0.75] | 0<br>[-0.60, 0.61] | 0.05<br>[-0.48, 0.58] | -0.57<sup>a</sup><br>[-1.19, 0.05] | 0.45 | 19 | 57 | 0.12, 0.52 |
|  | RF | **14.68**<br>[14.27, 15.10] | -0.06<br>[-0.53, 0.41] | -0.33<br>[-0.74, 0.08] | 0.06<br>[-0.30, 0.42] | -0.03<br>[-0.35, 0.29] | **-0.42<sup>b</sup>**<br>**[-0.79, -0.05]** | 0.45 | 19 | 57 | 0.16, 0.54 |
|  | OC | **2.21**<br>[2.08, 2.34] | -0.13<sup>a</sup><br>[-0.28, 0.02] | 0.13<sup>a</sup><br>[-0.01, 0.26] | -0.04<br>[-0.16, 0.09] | -0.07<br>[-0.17, 0.04] | -0.12<sup>a</sup><br>[-0.25, 0.01] | 0.37 | 19 | 57 | 0.32, 0.57 |
|  | RR | **4.22**<br>[4.00, 4.44] | 0.08<br>[-0.17, 0.33] | -0.06<br>[-0.28, 0.15] | 0.02<br>[-0.17, 0.21] | -0.06<br>[-0.23, 0.10] | **-0.27<sup>c</sup>**<br>**[-0.46, -0.08]** | 0.48 | 19 | 57 | 0.20, 0.58 |
| Species richness (Jackknife estimate) | All | **62.12**<br>[56.45, 67.79] | 3.44<br>[-2.77, 9.65] | -2.02<br>[-7.01, 2.96] | -2.31<br>[-6.36, 1.75] | -0.38<br>[-3.98, 3.23] | 3.73<sup>a</sup><br>[-0.31, 7.78] | 0.61 | 19 | 57 | 0.14, 0.67 |
|  | RF | **50.52**<br>[46.64, 54.40] | **5.83<sup>b</sup>**<br>[1.41, 10.26] | -2.47<br>[-6.33, 1.39] | -1.73<br>[-5.18, 1.72] | 0.07<br>[-2.96, 3.11] | 2.21<br>[-1.31, 5.73] | 0.44 | 19 | 57 | 0.35, 0.63 |
|  | OC | **10.98<sup>c</sup>**<br>[9.00, 12.97] | -1.58<br>[-3.80, 0.65] | **2.54<sup>c</sup>**<br>[0.65, 4.43] | -0.14<br>[-1.76, 1.48] | 0.29<br>[-1.15, 1.72] | 0.8<br>[-0.84, 2.44] | 0.51 | 19 | 57 | 0.35, 0.68 |
|  | RR | **11.58**<br>[10.46, 12.71] | **1.42<sup>b</sup>**<br>[0.15, 2.70] | -0.49<br>[-1.59, 0.61] | -0.85<sup>a</sup><br>[-1.81, 0.12] | -0.31<br>[-1.15, 0.54] | 0.84<sup>a</sup><br>[-0.14, 1.82] | 0.47 | 19 | 57 | 0.32, 0.64 |
| Density (birds, ha) | All | **42.68**<br>[38.51, 46.85] | **-8.61<sup>c</sup>**<br>[-13.36, -3.87] | -0.19<br>[-3.98, 3.60] | -1.11<br>[-4.26, 2.05] | -2.38<sup>a</sup><br>[-5.19, 0.43] | **-3.68<sup>b</sup>**<br>**[-6.74, -0.63]** | 0.65 | 19 | 48 | 0.41, 0.79 |
|  | RF | **33.53**<br>[29.93, 37.13] | -2.18<br>[-6.54, 2.18] | -1.32<br>[-5.16, 2.52] | -0.5<br>[-4.25, 3.26] | -1.02<br>[-4.32, 2.27] | **-5.09<sup>b</sup>**<br>**[-8.88, -1.29]** | 0.37 | 19 | 48 | 0.23, 0.52 |
|  | OC* | **11.86**<br>[10.40, 13.33] | **-3.42<sup>c</sup>**<br>[-5.37, -1.47] | **3.07<sup>c</sup>**<br>[1.38, 4.76] | 0.77<br>[-1.49, 3.02] | 0.10<br>[-1.77, 1.97] | -0.19<br>[-2.60, 2.21] | NA | NA | 48 | 0.53, NA |
|  | RR | **11.42**<br>[9.88, 12.96] | 0.04<br>[-1.85, 1.93] | -0.84<br>[-2.52, 0.84] | -0.6<br>[-2.34, 1.13] | -0.19<br>[-1.71, 1.32] | -0.55<br>[-2.33, 1.23] | 0.29 | 19 | 48 | 0.09, 0.35 |
| | | (Intercept) | Area (log-transformed) | $\Delta$ PC 1 scores | $\Delta$ PC 2 scores | $\Delta$ PC 3 scores | $\Delta$ Season (time diff) | ICC | N | Obs. | $mR^2$, $cR^2$ |
| Compositional dissimilarity (0 - 1) | All | **0.32**<br>[0.29, 0.34] | **-0.07<sup>c</sup>**<br>[-0.10, -0.05] | 0.01<br>[-0.00, 0.02] | 0.01<sup>a</sup><br>[-0.00, 0.03] | **0.02<sup>c</sup>**<br>[0.01, 0.03] | **0.04<sup>c</sup>**<br>[0.03, 0.05] | 0.8 | 14 | 126 | 0.73, 0.95 |
|  | RF | **0.29**<br>[0.25, 0.32] | **-0.08<sup>c</sup>**<br>[-0.11, -0.06] | 0.00<br>[-0.01, 0.01] | **0.01<sup>b</sup>**<br>[0.00, 0.03] | **0.02<sup>c</sup>**<br>[0.01, 0.03] | **0.03<sup>c</sup>**<br>[0.02, 0.03] | 0.88 | 14 | 126 | 0.68, 0.96 |
|  | OC | **0.44**<br>[0.33, 0.55] | 0.03<br>[-0.06, 0.11] | **0.05<sup>c</sup>**<br>[0.02, 0.09] | **-0.07<sup>c</sup>**<br>[-0.11, -0.02] | **0.06<sup>c</sup>**<br>[0.02, 0.10] | **0.13<sup>c</sup>**<br>[0.10, 0.16] | 0.85 | 14 | 126 | 0.26, 0.89 |
|  | RR | **0.23**<br>[0.20, 0.27] | **-0.06<sup>c</sup>**<br>[-0.09, -0.04] | 0.00<br>[-0.01, 0.01] | **0.03<sup>c</sup>**<br>[0.01, 0.04] | **0.03<sup>c</sup>**<br>[0.02, 0.04] | **0.03<sup>c</sup>**<br>[0.02, 0.04] | 0.79 | 14 | 126 | 0.60, 0.93 |

\*Mixed model did not converge, and we used a simple linear model without fragment identity as a random model instead

**Figure 2:**
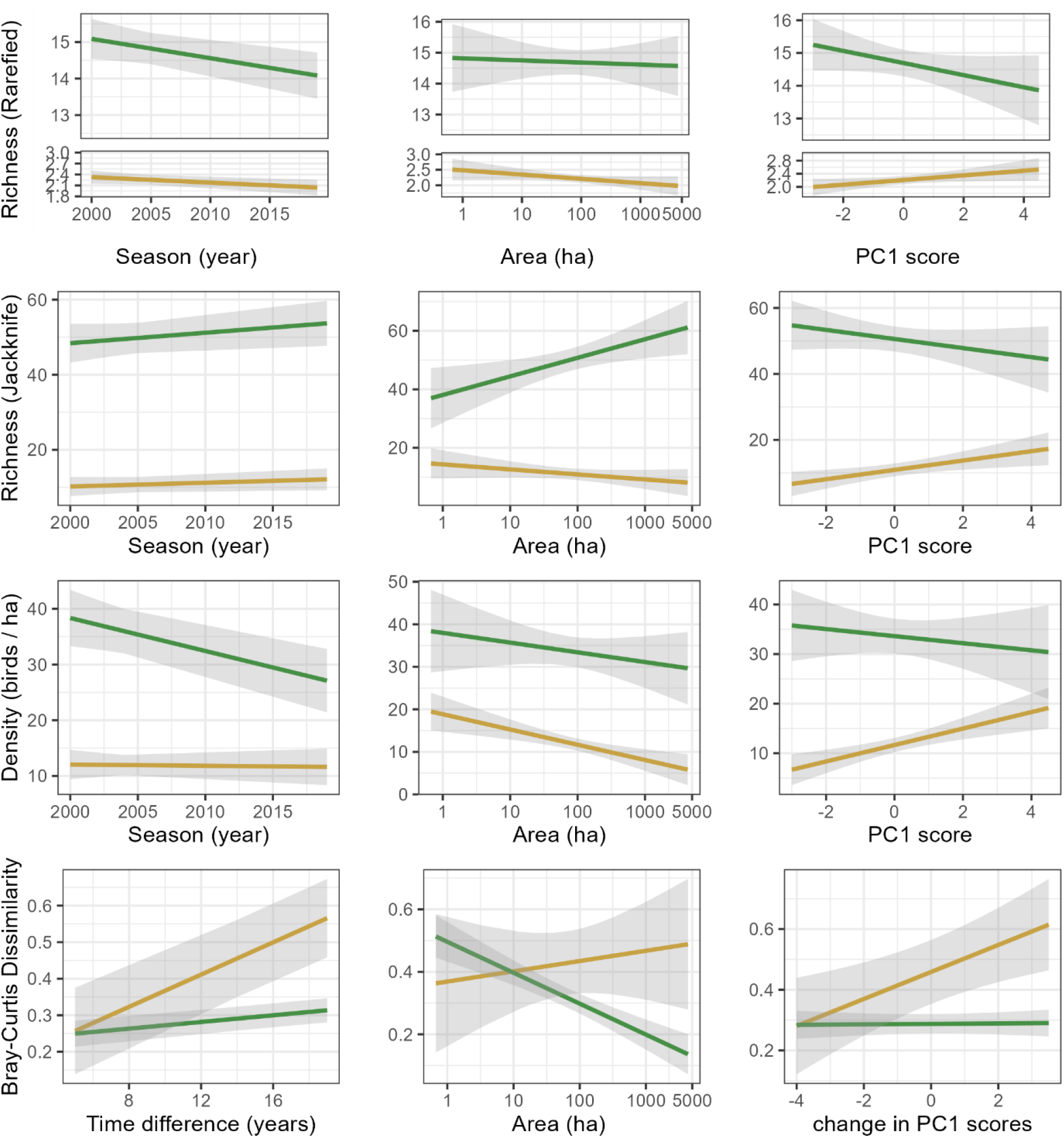
Effect plots showing the relationship between bird community variables and time, fragment area, and habitat structure (represented by the first principal component score, PC1). Curves are plotted separately for rainforest birds (RF, green) and open-country birds (OC, yellow). In models with compositional dissimilarity as the response, we used change in PC scores (ΔPC) over time and time difference between two sampling windows as predictors. The grey shaded area indicates 95% confidence intervals. See Table S2 for more information of each of the PC axes and the loadings of different habitat variables, and Table 2 for model summaries.

### Birds in relation to fragment size and habitat characteristics

When we statistically controlled time and habitat characteristics, rarefied richness and density of both rainforest and range-restricted birds did not change with fragment size (P > 0.05); rarefied richness of open-country birds was agnostic to fragment size (P > 0.05) but their density declined with area (*P* < 0.01) and the largest fragment in our landscape had 70% fewer open-country birds than the smallest (Figure 2, Table 2). Overall rarefied richness did not change, but the overall density of birds declined by a moderate 6% across three orders of magnitude in fragment size (*P* < 0.01).

In contrast to the rarefaction estimates, jackknife richness of rainforest and range-restricted birds in the largest fragment were 65% (*P* = 0.01) and 71% (*P* = 0.03) higher than the smallest fragments, respectively; for both these two guilds, larger fragments also had lower compositional dissimilarity (or greater stability) (*P* < 0.01). Jackknife richness and compositional dissimilarity of open country birds did not change substantially with fragment size (*P* > 0.1).

The first PC axis of habitat variation (and change in PC1 scores through time) was positively associated with open-country birds’ density and jackknife richness (*P* < 0.01) and marginally associated with higher rarefied richness (*P* = 0.06). Larger changes in PC1 scores (ΔPC) were associated with greater compositional change in open-country birds (*P* < 0.01, Figure 2, Table 2). Community properties of rainforest and range-restricted birds (as well as the overall bird community) were not influenced by PC1 scores.

Density and richness across guilds were not influenced by PC2 scores (except for marginally higher jackknife richness of range-restricted birds in fragments with higher PC2 scores, *P* = 0.08). Larger ΔPC2 scores were associated with greater dissimilarity in the composition of rainforest (*P* < 0.01) and range-restricted birds (*P* < 0.01) as well as the overall bird community (marginal, *P* = 0.09), but with less dissimilarity in open-country birds (*P* < 0.01). Similarly, changes in PC3 scores were not associated with a change in density or richness, but larger ΔPC3 scores were associated with the greater compositional dissimilarity of all guilds (*P* < 0.01, Table 2).

Fragment identity was more important than all other predictors, as evident from the difference between conditional and marginal R^2^ for all the models. Predictors explained an average of 34.2% (8.8% to 73.4%) of the variance in each model in Table 2.

### Bird abundance thresholds in relation to area

By analysing point-count frequency (as proxy for abundance) using Threshold Indicator Taxa Analysis (TITAN), we identified abundance thresholds for 38 species in relation to fragment area. Other species had scores of either “purity” (proportion of bootstrapped replicates where the threshold direction matches the observed response, i.e., purely consistent response direction) or “reliability” (proportion of bootstrapped replicates where the threshold magnitude’s corresponding Indicator Values with a consistently statistically discernible *P*-value, i.e., reliably significant Indicator Value scores) less than 90%. The mean area threshold was 96.7 ha for species that declined in occurrence frequency with fragment area and 34.3 ha for species that increased in occurrence frequency with fragment area (Figure 3, Table S4).

**Figure 3:**
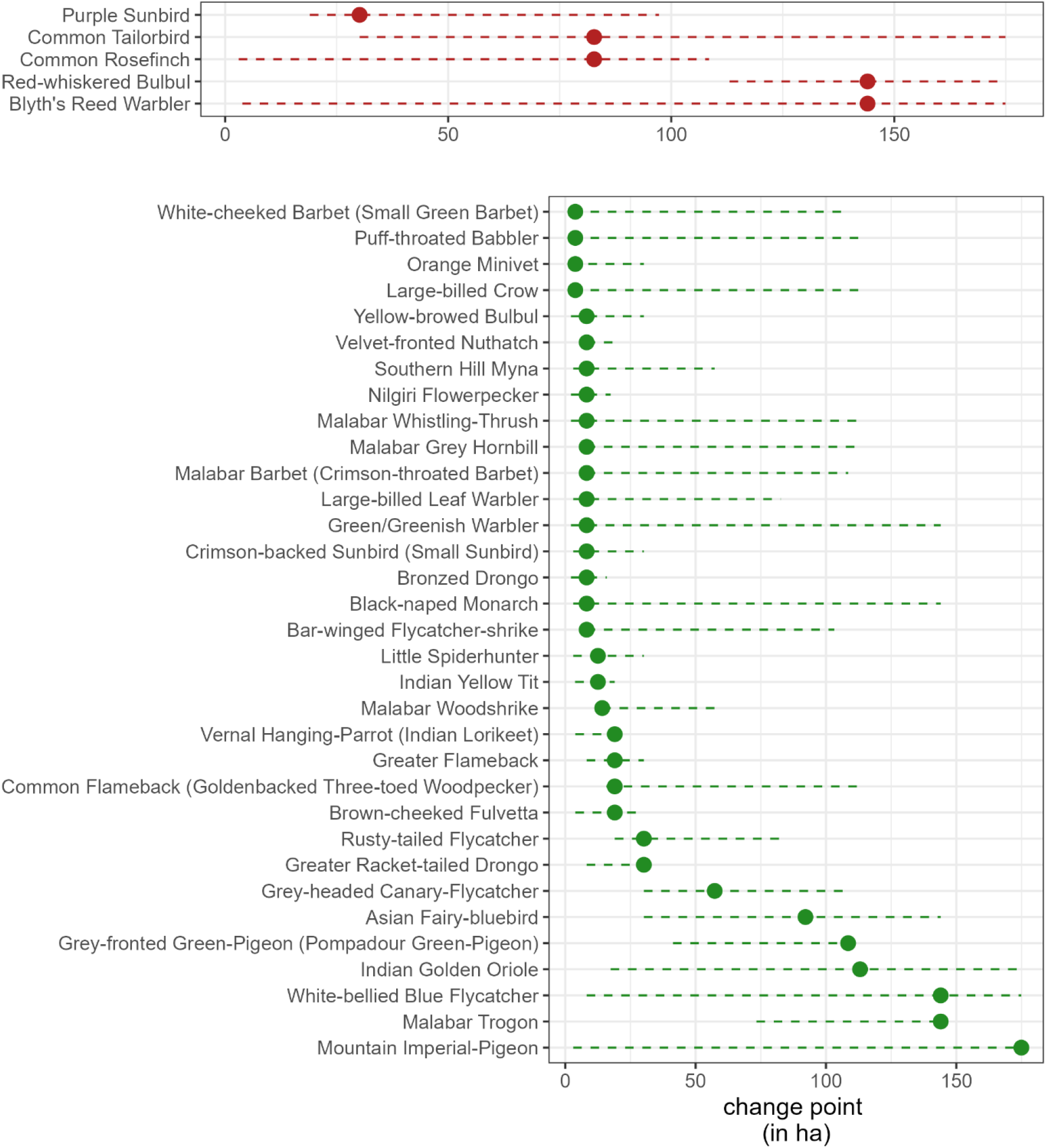
Thresholds in fragment area where bird occurrence frequency (a proxy for abundance) increases with area (species in green, increasers) or declines with area (red, decreasers) across small and medium-sized fragments (<300 ha). We used the Threshold Indicator Taxa Analysis (TITAN) algorithm to identify species-specific thresholds against the area gradient, used a 95% threshold in “purity” (proportion of bootstrapped replicates where the threshold direction matches the observed response, i.e. purely consistent response direction) and “reliability” (proportion of bootstrapped replicates where the threshold magnitude’s corresponding Indicator Values with a consistently statistically discernible P-value, i.e. reliably large Indicator Value scores) to subset relevant indicator taxon and plot the estimated thresholds and bootstrapped 95% confidence intervals.

Most bird species (33 of 38 species) with significant fragment size thresholds were positively associated with fragment size – i.e., increasers whose frequency was higher in larger than in smaller fragments (Figure 3, Table S4). All these species were rainforest specialists, and only seven of these species had area thresholds over 50 ha: Grey-headed Canary-flycatcher (threshold area: 57 ha), Asian Fairy-bluebird (92 ha), Grey-fronted Green-pigeon (108 ha), Indian Golden Oriole (113 ha), Malabar Trogon (144 ha), and White-bellied Blue-flycatcher (144 ha) and Mountain Imperial Pigeon (175 ha). In contrast, the five species that declined discernibly over fragment size thresholds – Purple Sunbird (in fragments larger than 30 ha), Common Rosefinch (83 ha), Common Tailorbird (83 ha), Blyth’s Reed Warbler (114 ha) and Red-whiskered Bulbul (144 ha) – were all open-habitat species (Figure 3, Table S4).

## Discussion

In long-isolated South Asian rainforest fragments, our study revealed an average decline of 11 rainforest birds per hectare per decade and a consequent decline of one species for every 23 birds: the community of open-country birds on the other hand remained relatively stable (Figure 2). Forest fragment area had a limited effect on measures of bird species richness and densities, with jackknife richness of rainforest and restricted-range birds increasing and densities overall and of open-country species decreasing with patch size (Table 2). Our study also identifies greater changes in species composition over wider time windows and with greater habitat structural change as indexed by ΔPC scores, with larger fragment areas tending to support more stable communities, particularly of rainforest and restricted-range birds (Table 2, Figure 2). These results provide a first look into decadal bird community dynamics in the rainforests of South Asia and corroborate observations on regional declines and interactive land-use and climate change impacts on rainforest birds from citizen science in the Western Ghats (SoIB 2020, Ramesh *et al.* 2022) as well as comprehensive bird community change elsewhere in the tropics and around the world (Şekercioğlu *et al.* 2019, Hendershot *et al.* 2020, Campos-Cerqueira *et al.* 2021, Tinoco *et al.* 2021, Williams & Fuente 2021).

Depressed survival and reduced breeding success drive reduction in bird populations in forest fragments, but persistent declines are probably linked to global or regional drivers of climate change. Increasing variability in precipitation can directly reduce nesting success (Brawn *et al.* 2017) while higher temperatures can directly reduce bird survival (Neate-Clegg *et al.* 2021), particularly in permontane and montane tropics where acclimatization via range-shifts are constrained by inter-specific competition (Srinivasan & Wilcove 2021). The effects may be indirect, too, where climate-driven declines in insect populations in turn constrain insectivorous birds (Raven & Wagner 2021). In southern India, birds breed in the dry season, when visibility is high, nest predators like snakes are less active and fledged birds can take advantage of heightened productivity during the monsoon rains (Ali *et al.* 1983, Shankar Raman 2002). However, regional changes in climate – increase in dry season rain due to stronger and more frequent cyclones (Balaguru *et al.* 2014) and decrease in monsoon rain due to rising surface temperatures further North (Varikoden *et al.* 2019) – are probably reducing populations of resident birds. Although follow-up studies are needed to identify the mechanism involved, our study reveals an overall pattern of decline.

Climate change can interact with land-use change in their impact on bird communities in forest fragments. Forest birds favour cooler, darker parts of the forest and because such habitats are uncommon in forest fragments, birds acclimatize by enhancing shuttling behaviour avoiding arid edge habitat (Tuff *et al.* 2016). Our study reveals a preferential effect of habitat quality and time period. Controlling for year of sampling, the richness of rainforest birds declines in fragments with poorer habitat quality (higher PC1 scores) but not their density, indicating some redundancy within rainforest birds. However, poorer habitat quality favoured density and richness (marginally so) of matrix-derived open-country birds, indicating lower redundancy (Figure 2). Citizen-science data corroborates this pattern at scale: poorer habitat ‘quality’ (or habitat complexity) correlates with lower occupancy of rainforest-adapted species (Ramesh *et al.* 2022). Rising temperatures can increase the fraction of edge habitat, pushing rainforest birds off their thermal optima while benefitting open-country birds. These effects may also account for the deviation from theoretical predictions from fragmentation – where persistent species loss long after fragmentation may not be lag-effects alone, but their interaction with climate.

Greater edge habitat (Ewers *et al.* 2007) and higher demographic stochasticity should render small fragments more vulnerable to decay in rainforest bird communities but landscape connectivity, especially long-distance dispersal over decadal scales, may buffer against any decay. Our study found that small fragments are just as vulnerable as large fragments: density and rarefied-richness of rainforest birds continued to decline when additive (Table 2) and interactive (Table S5) effects of fragment size are controlled for. However, consistent with global literature, we recorded higher overall richness (jackknife estimate) of rainforest birds in larger fragments, although not when controlling for variable densities of birds across fragments (rarefaction estimate), possibly indicating that rare bird species were more frequent in larger fragments and were represented better in the former estimate. We also found that large fragments had more stable communities of rainforest birds over time; moreover, stability was disproportionately high in large fragments (interactive models, Table S6). However, large fragments are critical for certain specialist and large-bodied bird species as indicated in our threshold analysis (Figure 3). This trend likely emerges from the traits of birds themselves – birds with large home-ranges or smaller home-range birds requiring better habitat structure – rather than guild-specific differences.

While rainforest bird density declined over time, the density of the subset of restricted-range birds in the aggregate did not show a significant decline. This contrasts with regional declines in endemic birds of the Western Ghats reported from citizen science studies (SoIB 2020), and landscape-level declines in species such as the endemic Malabar Grey Hornbill in the Anamalai Hills (Hariharan & Raman 2022). Stability in density of range-restricted species over time may be due to range-restricted species being limited in their geographic extent but not necessarily in abundance within specific parts of their range: Half of all range-restricted birds (11 of 22), such as White-cheeked Barbet *Psilopogon viridis*, Black-throated Munia *Lonchura kelaarti* and Malabar Barbet *Psilopogon malabaricus*, are commonly found in human modified landscapes like coffee plantations (Karanth *et al.* 2016), home gardens (Sidhu *et al.* 2010) and urban wooded areas (Praveen *et al.* 2022) within their restricted geographic ranges. The lack of decline in density aggregated across all RR species may hide declines in individual species, especially of more forest-restricted birds such as White-bellied Blue-Flycatcher, Nilgiri Thrush, White-bellied Sholakili, and Nilgiri Wood-pigeon (Table 2) (Warudkar *et al.* 2022).

Overall, we highlight a worrying decline in populations of rainforest birds, with potentially adverse impacts on bird conservation and downstream ecosystem functions in the future. Long-term conservation of tropical birds in old forest fragments therefore requires additional efforts such as protecting forest fragments irrespective of size (Turner & Corlett 1996), improving habitat quality through active and passive restoration (Hariharan & Raman 2022), and mitigating forest loss and fragmentation (Taubert *et al.* 2018). Maintaining ecosystem functions and stable populations of rainforest birds may further require mitigating regional stressors from global climate (Barlow *et al.* 2018, Lees *et al.* 2022, Ramesh *et al.* 2022).

## Acknowledgments

We thank the Science and Engineering Research Board, India (SERB, Project: EMR/2016/007968), Rohini Nilekani Philanthropies, and Arvind Datar for funding support. We thank the Tamil Nadu State Forest Department for permitting the research (Permit 74/2016), including V. K. Melkani, V. Ganesan, T. Paneerselvam, A. Xavier, and Krishnasamy, FD, DFOs and Range Officers of ATR. We are grateful to the management of Parry Agro Industries Ltd, Tata Coffee Ltd, and Tea Estates India Ltd for permission to survey sites within the estates. We are grateful to T. Vanidas, Manickraj, A. Sathish, T. Sundarraj, R. Rajesh and G. Moorthi for assistance with fieldwork. We would also like to thank Divya Mudappa, Anand M. Osuri, Hari Sridhar, Abhishek T. Gopal, Chayant Gonsalves, Swati Sidhu and P. Jeganathan for help in bird detectability classification and insightful discussions, and Hari Sridhar for sharing 2005 data.

