## Supplementary Material for "Forest bird decline and community change over 19 years in long-isolated South Asian tropical rainforest fragments"

**Supplementary Information**

A. **Extended methods**

*Why did we use an identity link for proportion and count data?*

We did not treat species richness and compositional dissimilarity as count and proportional data as is usually the case because our goal was to explain the difference between models rather than predict future community structure. Both richness and composition models passed all tests of model diagnostics. Moreover, rarefaction results in fractional richness, but Poisson-link count models require whole numbers: rounding off these rarefied estimates results in the loss of valuable ecological insight from differences in data. Similarly, compositional dissimilarity was not extreme enough (near zero or one) to warrant using models appropriate for proportional data.

*How did we incorporate bird species loudness and sightability covariates into distance sampling density estimation analysis?*

These two covariates were derived from scores made by nine experienced birders familiar with the regional avifauna, including the present authors. Each birder scored every bird species in the landscape on loudness (soft, moderate, loud) and ‘sightability’ (difficult, moderate, easy): the average score for each species was taken across all birders and grouped into thirds based on quantiles (see Table S1 for a list of bird taxa, habitat guilds, and corresponding detectability scores). The resulting groups on the ‘loudness’ and ‘sightability’ scale were used as covariates to model detectability using either half-normal or uniform detection functions with or without cosine adjustment terms. The estimated detection probability was used to estimate abundance and scaled by sampling effort to obtain density. In some cases, we did not detect sufficient individuals within a fragment to evaluate open-country birds (e.g., Akkamalai-Iyerpadi in the year 2002); in such cases, we manually added a near-zero estimate of 0.01 birds/ha

B. Tables

Table S1. Average sightability scores, loudness scores and number of detections of rainforest (RF), restricted-range (RR) and open-country (OC) birds in this study. The ID level column indicates confidence level of identification, at the species (S, 99.15% of detections), genus (G, 0.32% of detections), broader taxonomic group (T, 0.27% of detections) or no other information beyond a bird (B, when no ID was possible in the field, 0.26%).

| **No.** | **Common Name** | **ID level** | **Guild** | **Restricted-Range** | **Loudness** | **Sightability** | **Number of detections** | **Scientific Name** |
| --- | --- | --- | --- | --- | --- | --- | --- | --- |
| 1 | *Accipiter* sp. | G | RF |  | moderate | moderate | 1 | *Accipiter* sp. |
| 2 | *Acrocephalus* sp. | G | RF |  | soft | difficult | 2 | *Acrocephalus* sp. |
| 3 | Ashy Drongo | S | RF |  | loud | easy | 131 | *Dicrurus leucophaeus* |
| 4 | Ashy Prinia | S | OC |  | moderate | moderate | 10 | *Prinia socialis* |
| 5 | Ashy Woodswallow | S | OC |  | soft | easy | 2 | *Artamus fuscus* |
| 6 | Ashy/Brown-rumped Minivet | S | RF |  | soft | difficult | 1 | *Pericrocotus divaricatus/cantonensis* |
| 7 | Asian Brown Flycatcher | S | OC |  | soft | difficult | 31 | *Muscicapa dauurica* |
| 8 | Asian Emerald Dove | S | RF |  | moderate | moderate | 100 | *Chalcophaps indica* |
| 9 | Asian Fairy-bluebird | S | RF |  | loud | easy | 364 | *Irena puella* |
| 10 | Asian Koel | S | OC |  | loud | easy | 5 | *Eudynamys scolopaceus* |
| 11 | Bar-winged Flycatcher-Shrike | S | RF |  | soft | moderate | 152 | *Hemipus picatus* |
| 12 | Besra | S | RF |  | moderate | difficult | 39 | *Accipiter virgatus* |
| 13 | Bird sp. | B | NA |  | soft | difficult | 57 | Aves sp. |
| 14 | Black-and-orange Flycatcher | S | RF | Yes | soft | moderate | 123 | *Ficedula nigrorufa* |
| 15 | Black-headed Cuckooshrike | S | OC |  | soft | moderate | 3 | *Lalage melanoptera* |
| 16 | Black-naped Monarch | S | RF |  | loud | moderate | 342 | *Hypothymis azurea* |
| 17 | Black-rumped Flameback | S | OC |  | loud | easy | 26 | *Dinopium benghalense* |
| 18 | Black-Throated Munia | S | RF | Yes | soft | moderate | 6 | *Lonchura kelaarti* |
| 19 | Black-Winged Kite | S | OC |  | soft | easy | 2 | *Elanus caeruleus* |
| 20 | Black Eagle | S | RF |  | soft | easy | 3 | *Ictinaetus malaiensis* |
| 21 | Blue-bearded Bee-eater | S | RF |  | moderate | moderate | 1 | *Nyctyornis athertoni* |
| 22 | Blue-capped Rock-Thrush | S | RF |  | soft | difficult | 3 | *Monticola cinclorhyncha* |
| 23 | Blue-throated Flycatcher | S | RF |  | soft | difficult | 1 | *Cyornis rubeculoides* |
| 24 | Blue Flycatcher sp. | G | RF |  | soft | difficult | 6 | *Cyornis* sp. |
| 25 | Blyth's Reed Warbler | S | OC |  | moderate | difficult | 1139 | *Acrocephalus dumetorum* |
| 26 | Booted Warbler | S | OC |  | soft | difficult | 3 | *Iduna caligata* |
| 27 | Bronzed Drongo | S | RF |  | loud | easy | 240 | *Dicrurus aeneus* |
| 28 | Brown-breasted Flycatcher | S | RF |  | soft | difficult | 24 | *Muscicapa muttui* |
| 29 | Brown-capped Pygmy Woodpecker | S | RF |  | moderate | moderate | 24 | *Yungipicus nanus* |
| 30 | Brown-cheeked Fulvetta | S | RF |  | loud | difficult | 623 | *Alcippe poioicephala* |
| 31 | Brown Fish-Owl | S | RF |  | soft | moderate | 5 | *Ketupa zeylonensis* |
| 32 | Brown Shrike | S | OC |  | moderate | easy | 34 | *Lanius cristatus* |
| 33 | Chestnut-headed Bee-Eater | S | RF |  | loud | easy | 70 | *Merops leschenaulti* |
| 34 | Chestnut-tailed/Malabar Starling | S | RF | Yes | moderate | moderate | 11 | *Sturnia malabarica/blythii* |
| 35 | Chestnut-winged Cuckoo | S | RF |  | soft | difficult | 1 | *Clamator coromandus* |
| 36 | Cinereous Tit | S | OC |  | moderate | moderate | 3 | *Parus cinereus* |
| 37 | Common Flameback | S | RF |  | moderate | moderate | 68 | *Dinopium javanense* |
| 38 | Common Hawk-Cuckoo | S | OC |  | loud | difficult | 3 | *Hierococcyx varius* |
| 39 | Common Iora | S | OC |  | moderate | difficult | 83 | *Aegithina tiphia* |
| 40 | Common Rosefinch | S | OC |  | soft | moderate | 23 | *Carpodacus erythrinus* |
| 41 | Common Tailorbird | S | OC |  | moderate | moderate | 101 | *Orthotomus sutorius* |
| 42 | Crested Hawk-Eagle | S | RF |  | loud | easy | 1 | *Nisaetus cirrhatus* |
| 43 | Crested Serpent-Eagle | S | RF |  | loud | easy | 10 | *Spilornis cheela* |
| 44 | Crimson-Backed Sunbird | S | RF | Yes | moderate | moderate | 1712 | *Leptocoma minima* |
| 45 | Dark-Fronted Babbler | S | RF |  | soft | difficult | 136 | *Rhopocichla atriceps* |
| 46 | Diurnal Raptor sp. | B | NA |  | soft | easy | 2 | Accipitriformes/Falconiformes sp. |
| 47 | Drongo sp. | G | NA |  | loud | easy | 12 | *Dicrurus* sp. |
| 48 | Eurasian Hoopoe | S | OC |  | moderate | moderate | 10 | *Upupa epops* |
| 49 | *Ficedula* sp. | G | RF |  | soft | difficult | 1 | *Ficedula* sp. |
| 50 | Flame-Throated Bulbul | S | RF | Yes | moderate | moderate | 149 | *Rubigula gularis* |
| 51 | Flameback sp. | G | NA |  | loud | moderate | 3 | *Dinopium* sp. |
| 52 | Forest Wagtail | S | RF |  | soft | difficult | 33 | *Dendronanthus indicus* |
| 53 | Fork-Tailed Drongo-Cuckoo | S | RF |  | loud | difficult | 2 | *Surniculus dicruroides* |
| 54 | Golden-Fronted Leafbird | S | RF |  | moderate | difficult | 47 | *Chloropsis aurifrons* |
| 55 | Great Hornbill | S | RF |  | loud | easy | 28 | *Buceros bicornis* |
| 56 | Greater Coucal | S | OC |  | loud | easy | 36 | *Centropus sinensis* |
| 57 | Greater Flameback | S | RF |  | loud | moderate | 161 | *Chrysocolaptes guttacristatus* |
| 58 | Greater Racket-Tailed Drongo | S | RF |  | loud | easy | 435 | *Dicrurus paradiseus* |
| 59 | Greater/Common Flameback | G | RF |  | loud | moderate | 2 | *Chrysocolaptes guttacristatus/Dinopium javanense* |
| 60 | Green/Greenish Warbler ^[$]^ | S | RF |  | moderate | difficult | 1383 | *Phylloscopus nitidus/trochiloides* |
| 61 | Grey-Bellied Cuckoo | S | OC |  | moderate | difficult | 2 | *Cacomantis passerinus* |
| 62 | Grey-Breasted Prinia | S | OC |  | moderate | moderate | 29 | *Prinia hodgsonii* |
| 63 | Grey-Fronted Green-Pigeon | S | RF | Yes | moderate | difficult | 67 | *Treron affinis* |
| 64 | Grey-Headed Bulbul | S | RF | Yes | moderate | moderate | 21 | *Brachypodius priocephalus* |
| 65 | Grey-Headed Canary-Flycatcher | S | RF |  | moderate | easy | 431 | *Culicicapa ceylonensis* |
| 66 | Grey Junglefowl | S | RF |  | loud | moderate | 106 | *Gallus sonneratii* |
| 67 | Grey Wagtail | S | RF |  | moderate | easy | 16 | *Motacilla cinerea* |
| 68 | Heart-Spotted Woodpecker | S | RF |  | moderate | moderate | 23 | *Hemicircus canente* |
| 69 | House Crow | S | OC |  | loud | easy | 4 | *Corvus splendens* |
| 70 | Indian Blackbird | S | RF |  | moderate | difficult | 78 | *Turdus simillimus* |
| 71 | Indian Blue Robin | S | RF |  | moderate | difficult | 246 | *Larvivora brunnea* |
| 72 | Indian Golden Oriole | S | OC |  | moderate | easy | 39 | *Oriolus kundoo* |
| 73 | Indian Paradise-Flycatcher | S | RF |  | moderate | easy | 120 | *Terpsiphone paradisi* |
| 74 | Indian Peafowl | S | OC |  | loud | easy | 3 | *Pavo cristatus* |
| 75 | Indian Pitta | S | RF |  | loud | difficult | 1 | *Pitta brachyura* |
| 76 | Indian Pond-Heron | S | OC |  | soft | easy | 4 | *Ardeola grayii* |
| 77 | Indian Scimitar-Babbler | S | RF |  | loud | moderate | 341 | *Pomatorhinus horsfieldii* |
| 78 | Indian Yellow Tit | S | RF |  | moderate | moderate | 176 | *Machlolophus aplonotus* |
| 79 | Jungle Babbler | S | OC |  | loud | easy | 6 | *Turdoides striata* |
| 80 | Jungle Myna | S | OC |  | loud | easy | 4 | *Acridotheres fuscus* |
| 81 | Jungle Owlet | S | RF |  | loud | difficult | 2 | *Glaucidium radiatum* |
| 82 | Large-Billed Crow | S | OC |  | loud | easy | 323 | *Corvus macrorhynchos* |
| 83 | Large-Billed Leaf Warbler | S | RF |  | loud | difficult | 770 | *Phylloscopus magnirostris* |
| 84 | Large Hawk-Cuckoo | S | RF |  | soft | difficult | 3 | *Hierococcyx sparverioides* |
| 85 | Lesser Yellownape | S | RF |  | moderate | moderate | 22 | *Picus chlorolophus* |
| 86 | Lesser/Grey-Headed Fish-Eagle | G | RF |  | loud | easy | 1 | *Haliaeetus humilis/ichthyaetus* |
| 87 | Little Spiderhunter | S | RF |  | moderate | moderate | 373 | *Arachnothera longirostra* |
| 88 | Malabar Barbet | S | RF | Yes | loud | moderate | 123 | *Psilopogon malabaricus* |
| 89 | Malabar Grey Hornbill | S | RF | Yes | loud | easy | 270 | *Ocyceros griseus* |
| 90 | Malabar Parakeet | S | RF | Yes | loud | easy | 185 | *Psittacula columboides* |
| 91 | Malabar Trogon | S | RF |  | moderate | difficult | 75 | *Harpactes fasciatus* |
| 92 | Malabar Whistling-Thrush | S | RF |  | loud | moderate | 544 | *Myophonus horsfieldii* |
| 93 | Malabar Woodshrike | S | RF | Yes | moderate | moderate | 135 | *Tephrodornis sylvicola* |
| 94 | Martin/Swallow sp. | T | OC |  | soft | easy | 2 | *Hirundinidae* sp. |
| 95 | Mountain Imperial-Pigeon | S | RF |  | loud | moderate | 243 | *Ducula badia* |
| 96 | *Muscicapa* sp. | G | RF |  | soft | difficult | 30 | *Muscicapa* sp. |
| 97 | Myna/Starling sp. | G | OC |  | moderate | easy | 1 | *Sturnidae* sp. |
| 98 | Nilgiri Flowerpecker | S | RF | Yes | moderate | moderate | 1961 | *Dicaeum concolor* |
| 99 | Nilgiri Flycatcher | S | RF | Yes | soft | difficult | 296 | *Eumyias albicaudatus* |
| 100 | Nilgiri Thrush | S | RF | Yes | soft | difficult | 3 | *Zoothera neilgherriensis* |
| 101 | Nilgiri Wood-Pigeon | S | RF | Yes | soft | difficult | 3 | *Columba elphinstonii* |
| 102 | *Nisaetus* sp. | G | RF |  | moderate | moderate | 1 | *Nisaetus* sp. |
| 103 | Old World Flycatcher/Chat sp. | T | RF |  | moderate | moderate | 9 | *Muscicapidae* sp. |
| 104 | Old World Oriole sp. | G | RF |  | moderate | easy | 6 | *Oriolus* sp. |
| 105 | Orange-Headed Thrush | S | RF |  | moderate | difficult | 241 | *Geokichla citrina* |
| 106 | Orange Minivet | S | RF |  | moderate | easy | 494 | *Pericrocotus flammeus* |
| 107 | Oriental Dollarbird | S | RF |  | moderate | easy | 4 | *Eurystomus orientalis* |
| 108 | Oriental Honey-Buzzard | S | RF |  | soft | easy | 4 | *Pernis ptilorhynchus* |
| 109 | Oriental Magpie-Robin | S | OC |  | loud | easy | 30 | *Copsychus saularis* |
| 110 | Oriental Scops-Owl | S | RF |  | moderate | difficult | 1 | *Otus sunia* |
| 111 | Oriental White-Eye | S | RF |  | soft | moderate | 904 | *Zosterops palpebrosus* |
| 112 | Owl sp. | T | NA |  | soft | difficult | 1 | *Strigiformes* sp. |
| 113 | Palani Laughingthrush | S | RF | Yes | moderate | moderate | 24 | *Montecincla fairbanki* |
| 114 | *Phylloscopus* sp. | G | RF |  | moderate | difficult | 9 | *Phylloscopus sp.* |
| 115 | Pied Bushchat | S | OC |  | moderate | easy | 1 | *Saxicola caprata* |
| 116 | Pied Thrush | S | RF |  | soft | difficult | 3 | *Geokichla wardii* |
| 117 | Pigeon/Dove sp. | T | RF |  | moderate | difficult | 1 | *Columbidae* sp. |
| 118 | Plum-Headed Parakeet | S | OC |  | loud | easy | 83 | *Psittacula cyanocephala* |
| 119 | Puff-Throated Babbler | S | RF |  | loud | difficult | 297 | *Pellorneum ruficeps* |
| 120 | Purple Sunbird | S | OC |  | moderate | moderate | 74 | *Cinnyris asiaticus* |
| 121 | Red-Wattled Lapwing | S | OC |  | loud | easy | 1 | *Vanellus indicus* |
| 122 | Red-Whiskered Bulbul | S | OC |  | loud | easy | 418 | *Pycnonotus jocosus* |
| 123 | Red Spurfowl | S | RF |  | loud | moderate | 56 | *Galloperdix spadicea* |
| 124 | Rufous Babbler | S | RF | Yes | loud | moderate | 34 | *Turdoides subrufa* |
| 125 | Rufous Woodpecker | S | RF |  | loud | moderate | 13 | *Micropternus brachyurus* |
| 126 | Rusty-Tailed Flycatcher | S | RF |  | soft | difficult | 225 | *Ficedula ruficauda* |
| 127 | Shikra | S | OC |  | moderate | moderate | 6 | *Accipiter badius* |
| 128 | Small Minivet | S | OC |  | moderate | moderate | 42 | *Pericrocotus cinnamomeus* |
| 129 | Southern Hill Myna | S | RF |  | loud | easy | 493 | *Gracula indica* |
| 130 | Speckled Piculet | S | RF |  | soft | difficult | 25 | *Picumnus innominatus* |
| 131 | Spot-Bellied Eagle-Owl | S | RF |  | soft | difficult | 1 | *Bubo nipalensis* |
| 132 | Spotted Dove | S | OC |  | moderate | easy | 24 | *Streptopelia chinensis* |
| 133 | Square-Tailed Bulbul | S | RF |  | loud | easy | 514 | *Hypsipetes ganeesa* |
| 134 | Sri Lanka Frogmouth | S | RF |  | moderate | difficult | 2 | *Batrachostomus moniliger* |
| 135 | Streak-Throated Woodpecker | S | OC |  | loud | moderate | 9 | *Picus xanthopygaeus* |
| 136 | Sunbird sp. | G | NA |  | loud | easy | 1 | *Nectariniidae* sp. |
| 137 | Thick-Billed Warbler | S | OC |  | moderate | difficult | 2 | *Arundinax aedon* |
| 138 | Thrush sp. | T | RF |  | soft | difficult | 16 | *Turdidae* sp. |
| 139 | Tickell's Leaf Warbler | S | RF |  | soft | difficult | 4 | *Phylloscopus affinis* |
| 140 | Tit sp. | G | NA |  | moderate | easy | 1 | *Paridae* sp. |
| 141 | Velvet-Fronted Nuthatch | S | RF |  | moderate | easy | 529 | *Sitta frontalis* |
| 142 | Verditer Flycatcher | S | RF |  | soft | moderate | 7 | *Eumyias thalassinus* |
| 143 | Vernal Hanging-Parrot | S | RF |  | loud | moderate | 355 | *Loriculus vernalis* |
| 144 | Waterfowl sp. | B | OC |  | soft | easy | 1 | *Anatidae* sp. |
| 145 | Western Crowned Warbler | S | RF |  | soft | difficult | 97 | *Phylloscopus occipitalis* |
| 146 | White-Bellied Blue Flycatcher | S | RF | Yes | soft | difficult | 196 | *Cyornis pallidipes* |
| 147 | White-Bellied Sholakili | S | RF | Yes | moderate | difficult | 45 | *Sholicola albiventris* |
| 148 | White-Bellied Treepie | S | RF | Yes | loud | easy | 69 | *Dendrocitta leucogastra* |
| 149 | White-Bellied Woodpecker | S | RF |  | loud | moderate | 16 | *Dryocopus javensis* |
| 150 | White-Breasted Waterhen | S | OC |  | loud | easy | 3 | *Amaurornis phoenicurus* |
| 151 | White-Cheeked Barbet | S | RF | Yes | loud | moderate | 652 | *Psilopogon viridis* |
| 152 | White-Throated Kingfisher | S | OC |  | loud | easy | 9 | *Halcyon smyrnensis* |
| 153 | Woodpecker sp. | T | RF |  | moderate | easy | 29 | *Picidae* sp. |
| 154 | Wynaad Laughingthrush | S | RF | Yes | moderate | difficult | 13 | *Ianthocincla delesserti* |
| 157 | Yellow-Browed Bulbul | S | RF |  | loud | easy | 1221 | *Iole indica* |

[$] – Previously considered sub-species, Green and Greenish Warblers were recorded as separate species only in the 2019 census; records were collapsed to allow comparison with previous censuses

**Table S2**: Principal Component (PC) Analysis of seven habitat variables indicating eigenvalues and variable loadings on each of the PC axes that represent orthogonal constituents. The component against which a variable is most strongly loaded is indicated in bold.

| Parameters / Principal Components | PC1 | PC2 | PC3 | PC4 | PC5 | PC6 | PC7 |
| --- | --- | --- | --- | --- | --- | --- | --- |
| Tree density (ha^-1^) | **-0.45** | 0.342 | -0.182 | 0.231 | -0.504 | -0.215 | -0.542 |
| Tree basal area (m² ha^-1^) | -0.463 | -0.095 | **0.466** | -0.224 | -0.333 | 0.616 | 0.137 |
| Tree species richness (rarefied*) | -0.333 | -0.342 | **-0.602** | 0.221 | 0.355 | 0.459 | -0.156 |
| Leaf litter depth (cm) | **-0.463** | -0.065 | 0.234 | 0.591 | 0.175 | -0.333 | 0.486 |
| Canopy cover (%) | **-0.385** | 0.354 | 0.217 | -0.405 | 0.656 | -0.134 | -0.259 |
| Vertical stratification (scored 0-8) | -0.327 | **-0.418** | -0.313 | -0.566 | -0.208 | -0.445 | 0.242 |
| Shrub density (m^-2^) | -0.052 | **0.673** | -0.433 | -0.118 | -0.067 | 0.192 | 0.55 |
| Eigenvalue | 2.395 | 1.499 | 0.982 | 0.819 | 0.573 | 0.393 | 0.339 |
| Variance % | 34.211 | 21.42 | 14.033 | 11.699 | 8.182 | 5.609 | 4.846 |
| Cumulative Variance % | 34.211 | 55.631 | 69.664 | 81.363 | 89.545 | 95.154 | 100 |

*Rarefaction estimate of tree species richness in a standardised sample of 29 individuals across all sites.

**Table S3**: Bird density and parameter estimates from point count survey distance sampling across fragments and seasons in the Anamalai Hills. Tabled values include bird density (individuals/ha), se – standard error, cv – coefficient of variation, lcl –lower 95% confidence limit, ucl – upper 95% confidence limit, df – degrees of freedom, AIC – Akaike Information Criterion, p – detection probability, distance bins in metres, and final selected model. RF – rainforest birds, OC – open-country birds, and RR – restricted range birds.

| **Season** | **Fragment** | **Guild** | **Density** | **se** | **cv** | **lcl** | **ucl** | **df** | **AIC** | **p** | **Distance bins** | **Model** |
| --- | --- | --- | --- | --- | --- | --- | --- | --- | --- | --- | --- | --- |
| 2000 | Akkamalai Iyerpadi | All | 38.0 | 2.67 | 0.07 | 33.10 | 43.63 | 5 | 1981.2 | 0.30 | [0,5,10,15,20,30,50] | half-normal |
| 2000 | Akkamalai Iyerpadi | OC | 2.7 | 0.90 | 0.34 | 1.31 | 5.49 | 3 | 196.1 | 0.49 | [0,10,20,30,50] | uniform |
| 2000 | Akkamalai Iyerpadi | RF | 37.0 | 2.65 | 0.07 | 32.17 | 42.62 | 5 | 1936.7 | 0.30 | [0,5,10,15,20,30,50] | half-normal |
| 2000 | Akkamalai Iyerpadi | RR | 15.4 | 2.10 | 0.14 | 11.77 | 20.12 | 2 | 682.9 | 0.22 | [0,5,10,15,20,30,50] | uniform |
| 2002 | Akkamalai Iyerpadi | All | 44.1 | 4.84 | 0.11 | 35.50 | 54.75 | 5 | 855.4 | 0.27 | [0,5,10,15,20,30,50] | half-normal |
| 2002 | Akkamalai Iyerpadi | OC | 0.01 | NA | NA | NA | NA | NA | NA | NA | NA | manual |
| 2002 | Akkamalai Iyerpadi | RF | 42.7 | 4.69 | 0.11 | 34.38 | 53.01 | 5 | 840.5 | 0.27 | [0,5,10,15,20,30,50] | half-normal |
| 2002 | Akkamalai Iyerpadi | RR | 13.8 | 1.70 | 0.12 | 10.76 | 17.58 | 1 | 270.6 | 0.25 | [0,5,10,15,20,30,50] | uniform |
| 2003 | Akkamalai Iyerpadi | All | 41.8 | 4.46 | 0.11 | 33.85 | 51.55 | 5 | 847.4 | 0.28 | [0,5,10,15,20,30,50] | half-normal |
| 2003 | Akkamalai Iyerpadi | OC | 0.7 | 0.47 | 0.62 | 0.00 | 1.67 |  | 95.0 | 1.00 | [0,10,20,30,50] | uniform |
| 2003 | Akkamalai Iyerpadi | RF | 40.4 | 4.37 | 0.11 | 32.60 | 49.96 | 5 | 829.0 | 0.29 | [0,5,10,15,20,30,50] | half-normal |
| 2003 | Akkamalai Iyerpadi | RR | 12.8 | 1.64 | 0.13 | 9.98 | 16.53 | 1 | 269.5 | 0.27 | [0,5,10,15,20,30,50] | uniform |
| 2004 | Akkamalai Iyerpadi | All | 51.3 | 3.50 | 0.07 | 44.88 | 58.70 | 5 | 2904.6 | 0.25 | [0,5,10,15,20,30,50] | half-normal |
| 2004 | Akkamalai Iyerpadi | OC | 5.0 | 1.32 | 0.26 | 1.90 | 26.45 | 4 | 413.1 | 0.20 | [0,5,10,15,20,30,50] | uniform |
| 2004 | Akkamalai Iyerpadi | RF | 49.0 | 3.46 | 0.07 | 42.69 | 56.34 | 5 | 2798.8 | 0.25 | [0,5,10,15,20,30,50] | half-normal |
| 2004 | Akkamalai Iyerpadi | RR | 22.5 | 3.55 | 0.16 | 16.55 | 30.66 | 3 | 888.8 | 0.16 | [0,5,10,15,20,30,50] | uniform |
| 2019 | Akkamalai Iyerpadi | All | 39.3 | 2.67 | 0.07 | 34.42 | 44.92 | 5 | 2897.6 | 0.34 | [0,5,10,15,20,30,50] | half-normal |
| 2019 | Akkamalai Iyerpadi | OC | 4.8 | 1.21 | 0.25 | 2.96 | 7.83 | 3 | 260.3 | 0.28 | [0,10,20,30,50] | uniform |
| 2019 | Akkamalai Iyerpadi | RF | 33.8 | 2.63 | 0.08 | 32.29 | 42.77 | 5 | 2755.8 | 0.35 | [0,5,10,15,20,30,50] | half-normal |
| 2019 | Akkamalai Iyerpadi | RR | 9.1 | 0.83 | 0.09 | 7.56 | 10.86 | 1 | 778.2 | 0.36 | [0,5,10,15,20,30,50] | uniform |
| 2000 | Andiparai | All | 45.2 | 5.55 | 0.12 | 35.49 | 57.64 | 5 | 1159.5 | 0.26 | [0,5,10,15,20,30,50] | half-normal |
| 2000 | Andiparai | OC | 2.1 | 1.04 | 0.49 | 0.83 | 5.39 | 3 | 205.4 | 0.60 | [0,5,10,15,20,30,50] | uniform |
| 2000 | Andiparai | RF | 44.8 | 5.52 | 0.12 | 35.17 | 57.19 | 5 | 1147.0 | 0.26 | [0,5,10,15,20,30,50] | half-normal |
| 2000 | Andiparai | RR | 10.4 | 1.26 | 0.12 | 8.19 | 13.21 | 1 | 350.2 | 0.30 | [0,5,10,15,20,30,50] | uniform |
| 2002 | Andiparai | All | 41.8 | 4.75 | 0.11 | 33.44 | 52.28 | 5 | 742.2 | 0.26 | [0,5,10,15,20,30,50] | half-normal |
| 2002 | Andiparai | OC | 3.4 | 1.47 | 0.43 | 1.79 | 13.90 | 4 | 136.1 | 0.26 | [0,5,10,15,20,30,50] | uniform |
| 2002 | Andiparai | RF | 41.0 | 4.72 | 0.12 | 32.72 | 51.45 | 5 | 727.5 | 0.25 | [0,5,10,15,20,30,50] | half-normal |
| 2002 | Andiparai | RR | 7.2 | 1.15 | 0.16 | 5.27 | 9.90 | 1 | 167.9 | 0.29 | [0,5,10,15,20,30,50] | uniform |
| 2019 | Andiparai | All | 34.1 | 3.10 | 0.09 | 28.57 | 40.80 | 5 | 1554.9 | 0.43 | [0,5,10,15,20,30,50] | half-normal |
| 2019 | Andiparai | OC | 8.3 | 5.62 | 0.68 | 2.44 | 28.25 | 4 | 258.8 | 0.17 | [0,5,10,15,20,30,50] | uniform |
| 2019 | Andiparai | RF | 31.6 | 2.97 | 0.09 | 26.25 | 37.97 | 5 | 1474.4 | 0.44 | [0,5,10,15,20,30,50] | half-normal |
| 2019 | Andiparai | RR | 9.3 | 2.68 | 0.29 | 5.30 | 16.25 | 3 | 316.0 | 0.25 | [0,5,10,15,20,30,50] | uniform |
| 2019 | Candura | All | 47.9 | 4.32 | 0.09 | 40.15 | 57.25 | 5 | 1752.8 | 0.28 | [0,5,10,15,20,30,50] | half-normal |
| 2019 | Candura | OC | 18.1 | 7.33 | 0.41 | 8.34 | 39.17 | 5 | 301.1 | 0.09 | [0,5,10,15,20,30,50] | uniform |
| 2019 | Candura | RF | 45.2 | 4.29 | 0.10 | 37.45 | 54.44 | 5 | 1602.7 | 0.27 | [0,5,10,15,20,30,50] | half-normal |
| 2019 | Candura | RR | 16.1 | 2.87 | 0.18 | 11.34 | 22.79 | 2 | 530.5 | 0.23 | [0,5,10,15,20,30,50] | uniform |
| 2002 | Injiparai | All | 42.7 | 4.62 | 0.11 | 34.48 | 52.83 | 5 | 868.9 | 0.28 | [0,5,10,15,20,30,50] | half-normal |
| 2002 | Injiparai | OC | 14.8 | 3.31 | 0.22 | 9.54 | 22.91 | 2 | 276.5 | 0.24 | [0,5,10,15,20,30,50] | uniform |
| 2002 | Injiparai | RF | 30.3 | 3.99 | 0.13 | 23.36 | 39.30 | 5 | 631.8 | 0.28 | [0,5,10,15,20,30,50] | half-normal |
| 2002 | Injiparai | RR | 14.8 | 3.87 | 0.26 | 8.89 | 24.65 | 2 | 232.0 | 0.19 | [0,5,10,15,20,30,50] | uniform |
| 2003 | Injiparai | All | 47.7 | 6.17 | 0.13 | 36.91 | 61.62 | 5 | 884.1 | 0.25 | [0,5,10,15,20,30,50] | half-normal |
| 2003 | Injiparai | OC | 23.5 | 8.68 | 0.37 | 11.58 | 47.79 | 3 | 257.1 | 0.14 | [0,5,10,15,20,30,50] | uniform |
| 2003 | Injiparai | RF | 34.1 | 4.10 | 0.12 | 26.91 | 43.17 | 5 | 661.3 | 0.26 | [0,5,10,15,20,30,50] | half-normal |
| 2003 | Injiparai | RR | 24.7 | 9.37 | 0.38 | 11.93 | 51.24 | 4 | 246.6 | 0.12 | [0,5,10,15,20,30,50] | uniform |
| 2004 | Injiparai | All | 37.1 | 3.66 | 0.10 | 30.56 | 45.05 | 5 | 930.8 | 0.35 | [0,5,10,15,20,30,50] | half-normal |
| 2004 | Injiparai | OC | 19.1 | 6.75 | 0.35 | 9.69 | 37.72 | 3 | 275.9 | 0.20 | [0,5,10,15,20,30,50] | uniform |
| 2004 | Injiparai | RF | 27.3 | 3.56 | 0.13 | 21.11 | 35.27 | 5 | 650.0 | 0.34 | [0,5,10,15,20,30,50] | half-normal |
| 2004 | Injiparai | RR | 10.4 | 1.77 | 0.17 | 7.39 | 14.50 | 1 | 252.9 | 0.33 | [0,5,10,15,20,30,50] | uniform |
| 2019 | Injiparai | All | 47.8 | 5.93 | 0.12 | 37.47 | 61.04 | 5 | 801.3 | 0.31 | [0,5,10,15,20,30,50] | half-normal |
| 2019 | Injiparai | OC | 10.8 | 2.99 | 0.28 | 6.24 | 18.70 | 1 | 181.9 | 0.27 | [0,5,10,15,20,30,50] | uniform |
| 2019 | Injiparai | RF | 38.8 | 5.35 | 0.14 | 29.62 | 50.87 | 5 | 650.1 | 0.32 | [0,5,10,15,20,30,50] | half-normal |
| 2019 | Injiparai | RR | 10.9 | 1.86 | 0.17 | 7.82 | 15.29 | 1 | 200.7 | 0.32 | [0,5,10,15,20,30,50] | uniform |
| 2019 | Iyerpadi Top | All | 37.5 | 3.35 | 0.09 | 31.46 | 44.73 | 5 | 1256.2 | 0.34 | [0,5,10,15,20,30,50] | half-normal |
| 2019 | Iyerpadi Top | OC | 22.1 | 5.84 | 0.26 | 13.24 | 36.99 | 3 | 336.7 | 0.12 | [0,5,10,15,20,30,50] | uniform |
| 2019 | Iyerpadi Top | RF | 24.4 | 2.58 | 0.11 | 19.79 | 29.98 | 5 | 965.4 | 0.41 | [0,5,10,15,20,30,50] | half-normal |
| 2019 | Iyerpadi Top | RR | 9.0 | 1.17 | 0.13 | 7.00 | 11.66 | 1 | 348.6 | 0.35 | [0,5,10,15,20,30,50] | uniform |
| 2000 | Karian Shola | All | 32.2 | 4.10 | 0.13 | 25.05 | 41.33 | 5 | 907.5 | 0.32 | [0,5,10,15,20,30,50] | half-normal |
| 2000 | Karian Shola | OC | 5.0 | 1.66 | 0.33 | 2.43 | 10.34 | 3 | 128.5 | 0.27 | [0,10,20,30,50] | uniform |
| 2000 | Karian Shola | RF | 32.5 | 4.64 | 0.14 | 24.58 | 43.00 | 5 | 858.6 | 0.30 | [0,5,10,15,20,30,50] | half-normal |
| 2000 | Karian Shola | RR | 11.1 | 1.53 | 0.14 | 8.45 | 14.57 | 1 | 337.8 | 0.31 | [0,5,10,15,20,30,50] | uniform |
| 2019 | Karian Shola | All | 13.8 | 2.47 | 0.18 | 9.73 | 19.55 | 5 | 884.8 | 0.67 | [0,5,10,15,20,30,50] | half-normal |
| 2019 | Karian Shola | OC | 5.6 | 2.48 | 0.44 | 2.42 | 12.95 | 4 | 290.1 | 0.26 | [0,5,10,15,20,30,50] | uniform |
| 2019 | Karian Shola | RF | 12.0 | 1.52 | 0.13 | 9.34 | 15.37 | 5 | 827.6 | 0.73 | [0,5,10,15,20,30,50] | half-normal |
| 2019 | Karian Shola | RR | 4.9 | 1.07 | 0.22 | 3.24 | 7.52 | 1 | 327.6 | 0.57 | [0,5,10,15,20,30,50] | uniform |
| 2000 | Korangamudi | All | 42.7 | 4.20 | 0.10 | 35.23 | 51.84 | 5 | 969.2 | 0.34 | [0,5,10,15,20,30,50] | half-normal |
| 2000 | Korangamudi | OC | 19.7 | 2.59 | 0.13 | 15.19 | 25.57 | 1 | 388.3 | 0.27 | [0,5,10,15,20,30,50] | uniform |
| 2000 | Korangamudi | RF | 23.6 | 3.47 | 0.15 | 17.70 | 31.58 | 5 | 595.7 | 0.39 | [0,5,10,15,20,30,50] | half-normal |
| 2000 | Korangamudi | RR | 3.8 | 1.06 | 0.28 | 2.22 | 6.54 | 1 | 161.4 | 0.69 | [0,5,10,15,20,30,50] | uniform |
| 2002 | Korangamudi | All | 46.8 | 4.88 | 0.10 | 38.05 | 57.46 | 5 | 926.5 | 0.27 | [0,5,10,15,20,30,50] | half-normal |
| 2002 | Korangamudi | OC | 15.9 | 2.40 | 0.15 | 11.78 | 21.48 | 1 | 314.0 | 0.25 | [0,5,10,15,20,30,50] | uniform |
| 2002 | Korangamudi | RF | 27.9 | 4.12 | 0.15 | 20.83 | 37.35 | 5 | 607.9 | 0.30 | [0,5,10,15,20,30,50] | half-normal |
| 2002 | Korangamudi | RR | 7.1 | 1.32 | 0.19 | 4.90 | 10.24 | 1 | 186.1 | 0.31 | [0,5,10,15,20,30,50] | uniform |
| 2004 | Korangamudi | All | 47.7 | 4.36 | 0.09 | 39.88 | 57.15 | 5 | 1315.4 | 0.25 | [0,5,10,15,20,30,50] | half-normal |
| 2004 | Korangamudi | OC | 31.9 | 8.30 | 0.26 | 19.24 | 52.85 | 3 | 455.6 | 0.12 | [0,5,10,15,20,30,50] | uniform |
| 2004 | Korangamudi | RF | 29.9 | 3.48 | 0.12 | 23.76 | 37.58 | 5 | 872.9 | 0.27 | [0,5,10,15,20,30,50] | half-normal |
| 2004 | Korangamudi | RR | 9.6 | 2.27 | 0.24 | 6.09 | 15.29 | 2 | 300.9 | 0.27 | [0,5,10,15,20,30,50] | uniform |
| 2019 | Korangamudi | All | 39.0 | 3.18 | 0.08 | 33.19 | 45.75 | 5 | 1518.1 | 0.34 | [0,5,10,15,20,30,50] | half-normal |
| 2019 | Korangamudi | OC | 13.9 | 2.71 | 0.19 | 9.50 | 20.35 | 2 | 386.6 | 0.20 | [0,5,10,15,20,30,50] | uniform |
| 2019 | Korangamudi | RF | 28.3 | 2.88 | 0.10 | 23.19 | 34.56 | 5 | 1146.6 | 0.37 | [0,5,10,15,20,30,50] | half-normal |
| 2019 | Korangamudi | RR | 7.5 | 1.00 | 0.13 | 5.75 | 9.73 | 1 | 351.8 | 0.37 | [0,5,10,15,20,30,50] | uniform |
| 2000 | Manamboly | All | 43.1 | 4.04 | 0.09 | 35.86 | 51.87 | 5 | 1373.0 | 0.32 | [0,5,10,15,20,30,50] | half-normal |
| 2000 | Manamboly | OC | 5.0 | 2.58 | 0.51 | 1.91 | 13.21 | 4 | 204.5 | 0.26 | [0,5,10,15,20,30,50] | uniform |
| 2000 | Manamboly | RF | 41.7 | 3.88 | 0.09 | 34.74 | 50.11 | 5 | 1335.1 | 0.32 | [0,5,10,15,20,30,50] | half-normal |
| 2000 | Manamboly | RR | 14.3 | 1.51 | 0.11 | 11.64 | 17.66 | 1 | 504.5 | 0.31 | [0,5,10,15,20,30,50] | uniform |
| 2002 | Manamboly | All | 51.6 | 7.12 | 0.14 | 39.24 | 67.92 | 5 | 802.6 | 0.24 | [0,5,10,15,20,30,50] | half-normal |
| 2002 | Manamboly | OC | 6.7 | 2.67 | 0.40 | 3.13 | 14.55 | 4 | 121.1 | 0.20 | [0,5,10,15,20,30,50] | uniform |
| 2002 | Manamboly | RF | 51.4 | 7.23 | 0.14 | 38.82 | 68.00 | 5 | 766.0 | 0.23 | [0,5,10,15,20,30,50] | half-normal |
| 2002 | Manamboly | RR | 17.7 | 2.20 | 0.12 | 13.82 | 22.65 | 1 | 297.3 | 0.24 | [0,5,10,15,20,30,50] | uniform |
| 2004 | Manamboly | All | 38.1 | 3.44 | 0.09 | 31.89 | 45.50 | 5 | 1232.9 | 0.29 | [0,5,10,15,20,30,50] | half-normal |
| 2004 | Manamboly | OC | 3.3 | 1.37 | 0.41 | 1.50 | 7.27 | 3 | 130.7 | 0.40 | [0,10,20,30,50] | uniform |
| 2004 | Manamboly | RF | 37.6 | 3.41 | 0.09 | 31.49 | 44.96 | 5 | 1198.8 | 0.29 | [0,5,10,15,20,30,50] | half-normal |
| 2004 | Manamboly | RR | 18.5 | 3.15 | 0.17 | 13.26 | 25.84 | 2 | 445.4 | 0.20 | [0,5,10,15,20,30,50] | uniform |
| 2019 | Manamboly | All | 15.6 | 1.21 | 0.08 | 13.38 | 18.18 | 5 | 1230.5 | 0.75 | [0,5,10,15,20,30,50] | half-normal |
| 2019 | Manamboly | OC | 18.6 | 11.54 | 0.62 | 6.00 | 57.62 | 5 | 259.7 | 0.09 | [0,5,10,15,20,30,50] | uniform |
| 2019 | Manamboly | RF | 25.3 | 3.18 | 0.13 | 19.80 | 32.39 | 5 | 1032.1 | 0.43 | [0,5,10,15,20,30,50] | half-normal |
| 2019 | Manamboly | RR | 11.4 | 1.53 | 0.13 | 8.76 | 14.83 | 1 | 460.9 | 0.36 | [0,5,10,15,20,30,50] | uniform |
| 2019 | Murugaali BlackBridge | All | 61.1 | 8.24 | 0.13 | 46.78 | 79.90 | 5 | 683.2 | 0.23 | [0,5,10,15,20,30,50] | half-normal |
| 2019 | Murugaali BlackBridge | OC | 19.6 | 5.58 | 0.29 | 11.17 | 34.28 | 2 | 176.0 | 0.16 | [0,5,10,15,20,30,50] | uniform |
| 2019 | Murugaali BlackBridge | RF | 44.9 | 6.77 | 0.15 | 33.32 | 60.46 | 5 | 526.8 | 0.24 | [0,5,10,15,20,30,50] | half-normal |
| 2019 | Murugaali BlackBridge | RR | 13.3 | 2.12 | 0.16 | 9.65 | 18.21 | 1 | 192.8 | 0.28 | [0,5,10,15,20,30,50] | uniform |
| 2019 | Murugaali Sholayar | All | 52.5 | 4.86 | 0.09 | 43.73 | 62.99 | 5 | 1677.5 | 0.28 | [0,5,10,15,20,30,50] | half-normal |
| 2019 | Murugaali Sholayar | OC | 19.2 | 4.82 | 0.25 | 11.81 | 31.34 | 3 | 363.5 | 0.13 | [0,5,10,15,20,30,50] | uniform |
| 2019 | Murugaali Sholayar | RF | 43.6 | 4.33 | 0.10 | 35.83 | 52.98 | 5 | 1405.9 | 0.29 | [0,5,10,15,20,30,50] | half-normal |
| 2019 | Murugaali Sholayar | RR | 14.0 | 1.53 | 0.11 | 11.27 | 17.35 | 1 | 492.6 | 0.29 | [0,5,10,15,20,30,50] | uniform |
| 2000 | OldValparai | All | 44.6 | 4.75 | 0.11 | 36.11 | 54.99 | 5 | 897.5 | 0.29 | [0,5,10,15,20,30,50] | half-normal |
| 2000 | OldValparai | OC | 15.7 | 8.11 | 0.52 | 5.91 | 41.58 | 3 | 164.9 | 0.12 | [0,5,10,15,20,30,50] | uniform |
| 2000 | OldValparai | RF | 39.3 | 4.02 | 0.10 | 32.11 | 48.06 | 5 | 804.1 | 0.29 | [0,5,10,15,20,30,50] | half-normal |
| 2000 | OldValparai | RR | 13.7 | 1.74 | 0.13 | 10.66 | 17.62 | 1 | 291.9 | 0.29 | [0,5,10,15,20,30,50] | uniform |
| 2002 | OldValparai | All | 39.7 | 4.94 | 0.12 | 31.08 | 50.76 | 5 | 795.1 | 0.28 | [0,5,10,15,20,30,50] | half-normal |
| 2002 | OldValparai | OC | 11.3 | 4.01 | 0.36 | 5.65 | 22.53 | 4 | 153.0 | 0.14 | [0,5,10,15,20,30,50] | uniform |
| 2002 | OldValparai | RF | 34.8 | 4.50 | 0.13 | 26.92 | 44.91 | 5 | 725.6 | 0.29 | [0,5,10,15,20,30,50] | half-normal |
| 2002 | OldValparai | RR | 11.1 | 1.60 | 0.14 | 8.36 | 14.75 | 1 | 245.6 | 0.30 | [0,5,10,15,20,30,50] | uniform |
| 2004 | OldValparai | All | 51.7 | 6.16 | 0.12 | 40.85 | 65.43 | 5 | 975.7 | 0.26 | [0,5,10,15,20,30,50] | half-normal |
| 2004 | OldValparai | OC | 25.5 | 9.41 | 0.37 | 12.51 | 52.10 | 5 | 175.5 | 0.08 | [0,5,10,15,20,30,50] | uniform |
| 2004 | OldValparai | RF | 46.2 | 6.26 | 0.14 | 35.35 | 60.41 | 5 | 866.9 | 0.25 | [0,5,10,15,20,30,50] | half-normal |
| 2004 | OldValparai | RR | 11.0 | 1.51 | 0.14 | 8.36 | 14.39 | 1 | 262.9 | 0.31 | [0,5,10,15,20,30,50] | uniform |
| 2019 | OldValparai | All | 41.1 | 3.68 | 0.09 | 34.50 | 49.02 | 5 | 1343.9 | 0.34 | [0,5,10,15,20,30,50] | half-normal |
| 2019 | OldValparai | OC | 9.6 | 2.04 | 0.21 | 6.36 | 14.59 | 2 | 300.2 | 0.24 | [0,5,10,15,20,30,50] | uniform |
| 2019 | OldValparai | RF | 32.9 | 3.47 | 0.11 | 26.80 | 40.51 | 5 | 1119.7 | 0.36 | [0,5,10,15,20,30,50] | half-normal |
| 2019 | OldValparai | RR | 12.2 | 1.60 | 0.13 | 9.40 | 15.77 | 1 | 426.3 | 0.33 | [0,5,10,15,20,30,50] | uniform |
| 2000 | Pannimade | All | 53.5 | 4.60 | 0.09 | 45.23 | 63.38 | 5 | 1076.0 | 0.29 | [0,5,10,15,20,30,50] | half-normal |
| 2000 | Pannimade | OC | 13.8 | 7.23 | 0.52 | 5.17 | 37.03 | 5 | 150.5 | 0.11 | [0,5,10,15,20,30,50] | uniform |
| 2000 | Pannimade | RF | 49.2 | 4.39 | 0.09 | 41.26 | 58.60 | 5 | 1003.4 | 0.29 | [0,5,10,15,20,30,50] | half-normal |
| 2000 | Pannimade | RR | 15.1 | 1.80 | 0.12 | 11.92 | 19.12 | 1 | 317.8 | 0.27 | [0,5,10,15,20,30,50] | uniform |
| 2019 | Pannimade | All | 46.0 | 3.34 | 0.07 | 39.89 | 53.05 | 5 | 1724.8 | 0.34 | [0,5,10,15,20,30,50] | half-normal |
| 2019 | Pannimade | OC | 15.9 | 4.00 | 0.25 | 9.76 | 25.97 | 3 | 305.4 | 0.14 | [0,5,10,15,20,30,50] | uniform |
| 2019 | Pannimade | RF | 35.8 | 3.16 | 0.09 | 30.09 | 42.56 | 5 | 1433.5 | 0.37 | [0,5,10,15,20,30,50] | half-normal |
| 2019 | Pannimade | RR | 11.1 | 1.33 | 0.12 | 8.79 | 14.08 | 1 | 432.9 | 0.32 | [0,5,10,15,20,30,50] | uniform |
| 2000 | Puduthottam | All | 36.7 | 4.23 | 0.12 | 29.28 | 46.10 | 5 | 875.5 | 0.36 | [0,5,10,15,20,30,50] | half-normal |
| 2000 | Puduthottam | OC | 19.4 | 9.27 | 0.48 | 7.81 | 48.05 | 5 | 176.9 | 0.11 | [0,5,10,15,20,30,50] | uniform |
| 2000 | Puduthottam | RF | 32.2 | 4.34 | 0.13 | 24.67 | 41.94 | 5 | 756.6 | 0.36 | [0,5,10,15,20,30,50] | half-normal |
| 2000 | Puduthottam | RR | 11.4 | 1.72 | 0.15 | 8.43 | 15.34 | 1 | 285.7 | 0.34 | [0,5,10,15,20,30,50] | uniform |
| 2002 | Puduthottam | All | 34.7 | 3.71 | 0.11 | 28.17 | 42.86 | 5 | 790.0 | 0.32 | [0,5,10,15,20,30,50] | half-normal |
| 2002 | Puduthottam | OC | 13.9 | 4.87 | 0.35 | 7.00 | 27.44 | 4 | 157.8 | 0.12 | [0,5,10,15,20,30,50] | uniform |
| 2002 | Puduthottam | RF | 29.9 | 3.62 | 0.12 | 23.60 | 37.97 | 5 | 688.7 | 0.33 | [0,5,10,15,20,30,50] | half-normal |
| 2002 | Puduthottam | RR | 9.7 | 1.56 | 0.16 | 7.03 | 13.30 | 1 | 221.8 | 0.30 | [0,5,10,15,20,30,50] | uniform |
| 2003 | Puduthottam | All | 34.9 | 5.80 | 0.17 | 25.07 | 48.47 | 5 | 692.7 | 0.29 | [0,5,10,15,20,30,50] | half-normal |
| 2003 | Puduthottam | OC | 12.2 | 3.93 | 0.32 | 6.48 | 22.89 | 3 | 160.0 | 0.13 | [0,5,10,15,20,30,50] | uniform |
| 2003 | Puduthottam | RF | 30.2 | 5.29 | 0.18 | 21.33 | 42.69 | 5 | 622.1 | 0.31 | [0,5,10,15,20,30,50] | half-normal |
| 2003 | Puduthottam | RR | 10.2 | 1.65 | 0.16 | 7.42 | 14.04 | 1 | 210.2 | 0.28 | [0,5,10,15,20,30,50] | uniform |
| 2004 | Puduthottam | All | 39.4 | 3.54 | 0.09 | 33.02 | 47.03 | 5 | 1200.9 | 0.28 | [0,5,10,15,20,30,50] | half-normal |
| 2004 | Puduthottam | OC | 18.2 | 5.44 | 0.30 | 10.16 | 32.48 | 4 | 262.2 | 0.10 | [0,5,10,15,20,30,50] | uniform |
| 2004 | Puduthottam | RF | 34.7 | 3.49 | 0.10 | 28.50 | 42.34 | 5 | 1058.7 | 0.28 | [0,5,10,15,20,30,50] | half-normal |
| 2004 | Puduthottam | RR | 10.8 | 1.23 | 0.11 | 8.61 | 13.52 | 1 | 354.0 | 0.28 | [0,5,10,15,20,30,50] | uniform |
| 2019 | Puduthottam | All | 22.0 | 12.63 | 0.57 | 7.69 | 62.84 | 5 | 977.0 | 0.45 | [0,5,10,15,20,30,50] | half-normal |
| 2019 | Puduthottam | OC | 12.7 | 6.08 | 0.48 | 5.17 | 31.37 | 5 | 233.3 | 0.11 | [0,5,10,15,20,30,50] | uniform |
| 2019 | Puduthottam | RF | 18.7 | 12.84 | 0.69 | 5.49 | 63.48 | 5 | 888.9 | 0.50 | [0,5,10,15,20,30,50] | half-normal |
| 2019 | Puduthottam | RR | 7.8 | 1.89 | 0.24 | 4.82 | 12.47 | 2 | 297.9 | 0.30 | [0,5,10,15,20,30,50] | uniform |
| 2002 | Sankarankudi | All | 35.2 | 4.43 | 0.13 | 27.52 | 45.10 | 5 | 690.1 | 0.35 | [0,5,10,15,20,30,50] | half-normal |
| 2002 | Sankarankudi | OC | 13.9 | 11.52 | 0.83 | 3.23 | 60.13 | 5 | 126.5 | 0.11 | [0,5,10,15,20,30,50] | uniform |
| 2002 | Sankarankudi | RF | 32.2 | 3.90 | 0.12 | 25.37 | 40.84 | 5 | 644.7 | 0.36 | [0,5,10,15,20,30,50] | half-normal |
| 2002 | Sankarankudi | RR | 11.7 | 1.63 | 0.14 | 8.92 | 15.48 | 1 | 209.3 | 0.28 | [0,5,10,15,20,30,50] | uniform |
| 2000 | Selaliparai1 | All | 66.0 | 19.16 | 0.29 | 36.46 | 119.36 | 5 | 197.7 | 0.17 | [0,5,10,15,20,30,50] | half-normal |
| 2000 | Selaliparai1 | OC | 12.3 | 3.56 | 0.29 | 6.64 | 22.73 | 1 | 63.8 | 0.25 | [0,5,10,15,20,30,50] | uniform |
| 2000 | Selaliparai1 | RF | 49.8 | 19.06 | 0.38 | 22.95 | 107.98 | 5 | 141.0 | 0.17 | [0,5,10,15,20,30,50] | half-normal |
| 2000 | Selaliparai1 | RR | 13.1 | 5.05 | 0.39 | 5.69 | 29.96 | 1 | 54.4 | 0.22 | [0,5,10,15,20,30,50] | uniform |
| 2019 | Selaliparai1 | All | 47.5 | 6.45 | 0.14 | 36.28 | 62.13 | 5 | 538.4 | 0.29 | [0,5,10,15,20,30,50] | half-normal |
| 2019 | Selaliparai1 | OC | 20.5 | 8.79 | 0.43 | 8.96 | 46.77 | 3 | 150.7 | 0.16 | [0,5,10,15,20,30,50] | uniform |
| 2019 | Selaliparai1 | RF | 38.6 | 5.53 | 0.14 | 29.12 | 51.27 | 5 | 437.5 | 0.29 | [0,5,10,15,20,30,50] | half-normal |
| 2019 | Selaliparai1 | RR | 11.1 | 4.53 | 0.41 | 5.05 | 24.58 | 2 | 104.2 | 0.22 | [0,5,10,15,20,30,50] | uniform |
| 2000 | Selaliparai2 | All | 55.9 | 26.23 | 0.47 | 22.73 | 137.64 | 5 | 139.8 | 0.26 | [0,5,10,15,20,30,50] | half-normal |
| 2000 | Selaliparai2 | OC | 24.5 | 6.22 | 0.25 | 14.44 | 41.69 | 1 | 75.3 | 0.30 | [0,5,10,15,20,30,50] | uniform |
| 2000 | Selaliparai2 | RF | 29.0 | 24.24 | 0.84 | 6.30 | 133.44 | 5 | 68.3 | 0.25 | [0,5,10,15,20,30,50] | half-normal |
| 2000 | Selaliparai2 | RR | 5.7 | 4.04 | 0.71 | 1.34 | 23.93 | 1 | 21.9 | 0.45 | [0,5,10,15,20,30,50] | uniform |
| 2019 | Selaliparai2 | All | 65.2 | 12.81 | 0.20 | 43.87 | 96.99 | 5 | 338.1 | 0.21 | [0,5,10,15,20,30,50] | half-normal |
| 2019 | Selaliparai2 | OC | 12.7 | 3.43 | 0.27 | 7.23 | 22.31 | 1 | 96.4 | 0.28 | [0,5,10,15,20,30,50] | uniform |
| 2019 | Selaliparai2 | RF | 49.5 | 10.78 | 0.22 | 32.05 | 76.61 | 5 | 259.4 | 0.21 | [0,5,10,15,20,30,50] | half-normal |
| 2019 | Selaliparai2 | RR | 10.0 | 2.26 | 0.23 | 6.32 | 15.77 | 1 | 79.0 | 0.29 | [0,5,10,15,20,30,50] | uniform |
| 2000 | Varagaliar | All | 33.3 | 4.91 | 0.15 | 24.88 | 44.51 | 5 | 779.1 | 0.37 | [0,5,10,15,20,30,50] | half-normal |
| 2000 | Varagaliar | OC | 1.3 | 0.81 | 0.65 | 0.38 | 4.12 | 3 | 81.7 | 1.02 | [0,10,20,30,50] | uniform |
| 2000 | Varagaliar | RR | 12.2 | 1.87 | 0.15 | 8.99 | 16.51 | 1 | 292.9 | 0.33 | [0,5,10,15,20,30,50] | uniform |
| 2019 | Varagaliar | All | 16.8 | 3.56 | 0.21 | 11.67 | 18.09 | 5 | 972.1 | 0.77 | [0,5,10,15,20,30,50] | half-normal |
| 2019 | Varagaliar | RF | 16.2 | 3.70 | 0.23 | 7.08 | 39.65 | 5 | 850.7 | 0.65 | [0,10,20,30,50] | half-normal |
| 2019 | Varagaliar | RR | 5.4 | 0.99 | 0.18 | 3.77 | 7.72 | 1 | 347.4 | 0.58 | [0,5,10,15,20,30,50] | uniform |
| 2000 | Varatuparai123 | All | 48.2 | 7.01 | 0.15 | 36.06 | 64.39 | 5 | 548.1 | 0.22 | [0,5,10,15,20,30,50] | half-normal |
| 2000 | Varatuparai123 | OC | 23.2 | 7.83 | 0.34 | 11.90 | 45.09 | 2 | 204.5 | 0.15 | [0,5,10,15,20,30,50] | uniform |
| 2000 | Varatuparai123 | RF | 27.1 | 4.55 | 0.17 | 19.49 | 37.80 | 5 | 356.2 | 0.27 | [0,5,10,15,20,30,50] | half-normal |
| 2000 | Varatuparai123 | RR | 9.1 | 1.57 | 0.17 | 6.46 | 12.84 | 1 | 139.4 | 0.31 | [0,5,10,15,20,30,50] | uniform |
| 2019 | Varatuparai123 | All | 32.2 | 7.30 | 0.23 | 4.51 | 229.78 | 5 | 469.3 | 0.29 | [0,5,10,15,20,30,50] | half-normal |
| 2019 | Varatuparai123 | OC | 11.5 | 2.84 | 0.25 | 7.02 | 18.89 | 1 | 188.8 | 0.25 | [0,5,10,15,20,30,50] | uniform |
| 2019 | Varatuparai123 | RF | 17.6 | 5.45 | 0.31 | 0.79 | 520.79 | 5 | 320.4 | 0.34 | [0,5,10,15,20,30,50] | half-normal |
| 2019 | Varatuparai123 | RR | 15.1 | 6.08 | 0.40 | 6.98 | 32.80 | 3 | 151.7 | 0.14 | [0,5,10,15,20,30,50] | uniform |
| 2000 | Varatuparai4 | All | 56.1 | 18.76 | 0.33 | 28.59 | 110.11 | 5 | 174.9 | 0.21 | [0,5,10,15,20,30,50] | half-normal |
| 2000 | Varatuparai4 | OC | 12.3 | 4.38 | 0.36 | 5.87 | 25.58 | 1 | 76.7 | 0.43 | [0,5,10,15,20,30,50] | uniform |
| 2000 | Varatuparai4 | RF | 42.6 | 16.12 | 0.38 | 19.98 | 90.66 | 5 | 100.2 | 0.15 | [0,5,10,15,20,30,50] | half-normal |
| 2000 | Varatuparai4 | RR | 7.9 | 3.15 | 0.40 | 3.34 | 18.61 | 1 | 43.5 | 0.30 | [0,5,10,15,20,30,50] | uniform |
| 2019 | Varatuparai4 | All | 34.3 | 5.30 | 0.15 | 25.27 | 46.46 | 5 | 345.5 | 0.28 | [0,5,10,15,20,30,50] | half-normal |
| 2019 | Varatuparai4 | OC | 14.7 | 5.05 | 0.34 | 7.51 | 28.83 | 2 | 115.5 | 0.19 | [0,5,10,15,20,30,50] | uniform |
| 2019 | Varatuparai4 | RF | 20.1 | 4.59 | 0.23 | 14.22 | 33.41 | 5 | 239.5 | 0.32 | [0,5,10,15,20,30,50] | half-normal |
| 2019 | Varatuparai4 | RR | 4.9 | 1.52 | 0.31 | 2.64 | 9.01 | 1 | 87.4 | 0.47 | [0,5,10,15,20,30,50] | uniform |

**Table S4**: Results of Threshold Indicator Taxa Analysis (TITAN) of bird abundance in relation to the gradient in rainforest fragment area across 15 small and medium sized fragments of area < 300 ha. Tabled values indicate change point (ha); frequency or the number of 43 fragment-year combinations where the species was recorded; direction of change (1 – decreasers, 2 – increasers); indicator value; 95% confidence interval of change point (columns 5% CP and 95% CP); purity or the proportion of bootstrapped replicates where the threshold direction matches the observed response, i.e., purely consistent response direction; and reliability or the proportion of bootstrapped replicates with probability *P* ≤ 0.05 of an equal or larger Indicator Values (i.e., reliably significant Indicator Value scores); and a filter variable to select species.

| Bird ID | Change Point (CP) (ha) | Frequency | Direction | 5% CP | 95% CP | Indicator Value | Purity | Reliability |
| --- | --- | --- | --- | --- | --- | --- | --- | --- |
| Asian Fairy-bluebird | 92.13 | 30 / 43 | 2 | 30.15 | 144.02 | 83.23 | 1.00 | 1.00 |
| Bar-winged Flycatcher-shrike | 8.19 | 28 / 43 | 2 | 8.19 | 103.21 | 68.34 | 0.95 | 0.91 |
| Black-naped Monarch | 8.19 | 33 / 43 | 2 | 3.04 | 144.02 | 83.75 | 1.00 | 1.00 |
| Blyth's Reed Warbler | 144.02 | 41 / 43 | 1 | 3.85 | 174.95 | 82.86 | 0.92 | 1.00 |
| Bronzed Drongo | 8.19 | 38 / 43 | 2 | 2.22 | 15.95 | 89.13 | 0.98 | 1.00 |
| Brown-cheeked Fulvetta | 18.98 | 34 / 43 | 2 | 3.85 | 30.15 | 87.98 | 1.00 | 1.00 |
| Common Flameback  (Goldenbacked Three-toed Woodpecker) | 18.98 | 24 / 43 | 2 | 15.75 | 113.08 | 76.38 | 1.00 | 1.00 |
| Crimson-backed Sunbird  (Small Sunbird) | 8.19 | 42 / 43 | 2 | 3.04 | 30.15 | 87.32 | 1.00 | 1.00 |
| Greater Flameback | 18.98 | 33 / 43 | 2 | 8.19 | 30.15 | 79.60 | 1.00 | 0.99 |
| Greater Racket-tailed Drongo | 30.15 | 33 / 43 | 2 | 8.19 | 30.15 | 85.95 | 1.00 | 1.00 |
| Green/Greenish Warbler | 8.19 | 43 / 43 | 2 | 2.22 | 144.02 | 79.75 | 0.95 | 1.00 |
| Grey-fronted Green-Pigeon  (Pompadour Green-Pigeon) | 108.50 | 22 / 43 | 2 | 41.31 | 113.08 | 81.89 | 1.00 | 1.00 |
| Grey-headed Canary-Flycatcher | 57.32 | 22 / 43 | 2 | 30.15 | 108.50 | 84.64 | 1.00 | 1.00 |
| Indian Golden Oriole | 113.08 | 14 / 43 | 2 | 17.38 | 174.95 | 48.06 | 1.00 | 0.91 |
| Indian Yellow Tit | 12.52 | 38 / 43 | 2 | 3.77 | 18.98 | 80.39 | 1.00 | 1.00 |
| Large-billed Crow | 3.85 | 37 / 43 | 2 | 3.04 | 113.08 | 87.49 | 0.95 | 0.98 |
| Large-billed Leaf Warbler | 8.19 | 38 / 43 | 2 | 3.04 | 82.73 | 84.57 | 1.00 | 1.00 |
| Little Spiderhunter | 12.52 | 37 / 43 | 2 | 3.04 | 30.15 | 88.98 | 1.00 | 1.00 |
| Malabar Barbet  (Crimson-throated Barbet) | 8.19 | 28 / 43 | 2 | 8.19 | 108.50 | 75.68 | 0.98 | 0.98 |
| Malabar Grey Hornbill | 8.199 | 37 / 43 | 2 | 8.19 | 113.08 | 81.71 | 1.00 | 1.00 |
| Malabar Trogon | 144.02 | 14 / 43 | 2 | 73.32 | 144.02 | 74.11 | 1.00 | 1.00 |
| Malabar Whistling-Thrush | 8.19 | 42 / 43 | 2 | 2.22 | 113.08 | 72.18 | 1.00 | 0.99 |
| Malabar Woodshrike | 14.16 | 32 / 43 | 2 | 14.16 | 57.32 | 82.66 | 0.99 | 1.00 |
| Mountain Imperial-Pigeon | 174.95 | 29 / 43 | 2 | 3.04 | 174.95 | 73.57 | 1.00 | 0.97 |
| Nilgiri Flowerpecker | 8.19 | 43 / 43 | 2 | 2.22 | 17.38 | 81.94 | 0.99 | 1.00 |
| Orange Minivet | 3.85 | 41 / 43 | 2 | 2.22 | 30.15 | 83.19 | 0.99 | 0.99 |
| Puff-throated Babbler | 3.85 | 41 / 43 | 2 | 3.04 | 113.08 | 85.27 | 0.98 | 0.92 |
| Red-whiskered Bulbul | 144.02 | 39 / 43 | 1 | 113.08 | 174.95 | 86.19 | 0.99 | 0.98 |
| Rusty-tailed Flycatcher | 30.15 | 31 / 43 | 2 | 18.98 | 82.73 | 93.16 | 1.00 | 1.00 |
| Southern Hill Myna | 8.19 | 36 / 43 | 2 | 3.04 | 57.32 | 85.98 | 1.00 | 1.00 |
| Velvet-fronted Nuthatch | 8.19 | 41 / 43 | 2 | 8.19 | 18.98 | 82.04 | 1.00 | 1.00 |
| Vernal Hanging-Parrot  (Indian Lorikeet) | 18.98 | 40 / 43 | 2 | 3.85 | 18.98 | 83.42 | 1.00 | 1.00 |
| White-bellied Blue Flycatcher | 144.02 | 26 / 43 | 2 | 8.19 | 174.95 | 85.01 | 1.00 | 0.99 |
| White-cheeked Barbet  (Small Green Barbet) | 3.85 | 41 / 43 | 2 | 3.04 | 108.19 | 84.11 | 1.00 | 1.00 |
| Yellow-browed Bulbul | 8.19 | 41 / 43 | 2 | 2.22 | 30.15 | 84.54 | 1.00 | 1.00 |
| Common Rosefinch | 82.73 | 13 / 43 | 1 | 3.04 | 108.50 | 44.88 | 1.00 | 0.93 |
| Common Tailorbird | 82.73 | 30 / 43 | 1 | 30.15 | 174.95 | 69.76 | 1.00 | 0.97 |
| Purple Sunbird | 30.15 | 12 / 43 | 1 | 18.98 | 97.28 | 48.77 | 1.00 | 0.99 |

**Table S5:** Linear mixed model regressions of bird species richness (rarefaction and jackknife estimates), bird density (distance sampling estimate), and compositional dissimilarity of bird communities (Bray-Curtis dissimilarity) of birds against time, fragment area, and habitat quality as indexed by Principal Component (PC) scores, and time x area interaction term. Fragment identity is the random effect and the intraclass correlation coefficient (ICC) is the variance fraction explained by fragment identity. Tabled values indicate regression coefficients (95% confidence interval), sample size of fixed effects (N = 19 or 14 fragments) and random levels (Observations = 57, 48 or 126), and marginal (mR²) and conditional R-squared (cR²). [All = entire bird community; RF = rainforest birds; RR = restricted-range birds; OC = open-country birds]

|  | **Richness (all)** | **(rf)** | **(rr)** | **(oc)** | **Density (all)** | | **(rf)** | **(rr)** | | **(oc)** | **Composition (all)** | **(rf)** | **(rr)** | | **(oc)** |
| --- | --- | --- | --- | --- | --- | --- | --- | --- | --- | --- | --- | --- | --- | --- | --- |
| *Predictors* | *Estimates* | *Estimates* | *Estimates* | *Estimates* | *Estimates* | | *Estimates* | *Estimates* | *Estimates* | | *Estimates* | *Estimates* | | *Estimates* | *Estimates* |
| (Intercept) | 23.39 (22.68 – 24.09) | 14.69 (14.27 – 15.11) | 4.23 (4.01 – 4.45) | 2.21 (2.09 – 2.34) | 42.61 (38.50 – 46.73) | | 33.40 (29.78 – 37.01) | 11.35 (9.79 – 12.92) | 11.92 (10.43 – 13.40) | | 0.31 (0.29 – 0.34) | 0.29 (0.25 – 0.32) | | 0.23 (0.20 – 0.27) | 0.43 (0.31 – 0.55) |
| area | -0.40 (-1.21 – 0.41) | -0.10 (-0.58 – 0.38) | 0.05 (-0.20 – 0.31) | -0.14 (-0.29 – 0.02) | -8.25 (-13.04 – -3.47) | | -1.61 (-6.02 – 2.79) | 0.30 (-1.63 – 2.22) | -3.47 (-5.45 – -1.50) | | -0.07 (-0.09 – -0.05) | -0.08 (-0.10 – -0.05) | | -0.06 (-0.09 – -0.04) | 0.05 (-0.05 – 0.14) |
| PC1 | -0.07 (-0.80 – 0.66) | -0.39 (-0.83 – 0.04) | -0.11 (-0.34 – 0.12) | 0.12 (-0.02 – 0.25) | 0.38 (-3.72 – 4.49) | | -0.20 (-4.22 – 3.81) | -0.37 (-2.14 – 1.40) | 2.93 (1.17 – 4.68) | |  |  | |  |  |
| PC2 | -0.06 (-0.68 – 0.56) | 0.03 (-0.34 – 0.40) | -0.01 (-0.20 – 0.18) | -0.04 (-0.17 – 0.08) | -0.75 (-4.06 – 2.57) | | 0.11 (-3.63 – 3.85) | -0.33 (-2.05 – 1.40) | 0.67 (-1.62 – 2.96) | |  |  | |  |  |
| PC3 | 0.02 (-0.52 – 0.56) | -0.04 (-0.36 – 0.28) | -0.07 (-0.24 – 0.09) | -0.07 (-0.18 – 0.04) | -2.21 (-5.08 – 0.67) | | -0.80 (-4.03 – 2.43) | -0.09 (-1.57 – 1.40) | 0.10 (-1.79 – 1.99) | |  |  | |  |  |
| Season | -0.52 (-1.14 – 0.10) | -0.40 (-0.77 – -0.02) | -0.25 (-0.44 – -0.06) | -0.11 (-0.24 – 0.02) | -3.88 (-7.02 – -0.73) | | -5.55 (-9.27 – -1.82) | -0.76 (-2.50 – 0.98) | -0.09 (-2.54 – 2.35) | |  |  | |  |  |
| area * Season | 0.19 (-0.18 – 0.57) | 0.10 (-0.13 – 0.32) | 0.07 (-0.04 – 0.18) | 0.02 (-0.05 – 0.10) | -0.69 (-2.63 – 1.24) | | -1.92 (-4.21 – 0.38) | -0.89 (-1.96 – 0.19) | 0.52 (-1.01 – 2.05) | |  |  | |  |  |
| dPC1 |  |  |  |  |  | |  |  |  | | 0.01 (-0.00 – 0.02) | 0.00 (-0.01 – 0.01) | | -0.00 (-0.02 – 0.01) | 0.07 (0.03 – 0.11) |
| dPC2 |  |  |  |  |  | |  |  |  | | 0.01 (-0.00 – 0.02) | 0.01 (0.00 – 0.03) | | 0.03 (0.01 – 0.04) | -0.08 (-0.13 – -0.04) |
| dPC3 |  |  |  |  |  | |  |  |  | | 0.02 (0.01 – 0.03) | 0.02 (0.01 – 0.04) | | 0.03 (0.02 – 0.04) | 0.07 (0.03 – 0.11) |
| timediff |  |  |  |  |  | |  |  |  | | 0.04 (0.03 – 0.05) | 0.03 (0.02 – 0.04) | | 0.03 (0.02 – 0.04) | 0.14 (0.11 – 0.17) |
| area * timediff |  |  |  |  |  | |  |  |  | | -0.01 (-0.01 – -0.00) | -0.01 (-0.01 – -0.00) | | 0.00 (-0.00 – 0.01) | -0.03 (-0.04 – -0.01) |
| σ^2^ | 1.74 | 0.63 | 0.16 | 0.08 | 34.78 | 52.28 | | 11.65 | | 25.50 | 0.00 | 0.00 | 0.00 | | 0.01 |
| τ_00_ | 1.52 _Fragment_ | 0.53 _Fragment_ | 0.15 _Fragment_ | 0.04 _Fragment_ | 59.48 _Fragment_ | 34.89 _Fragment_ | | 5.78 _Fragment_ | | 0.00 _Fragment_ | 0.00 _Fragment_ | 0.00 _Fragment_ | 0.00 _Fragment_ | | 0.05 _Fragment_ |
| ICC | 0.47 | 0.46 | 0.48 | 0.35 | 0.63 | 0.40 | | 0.33 | |  | 0.81 | 0.87 | 0.79 | | 0.90 |
| N | 19 _Fragment_ | 19 _Fragment_ | 19 _Fragment_ | 19 _Fragment_ | 19 _Fragment_ | 19 _Fragment_ | | 19 _Fragment_ | | 19 _Fragment_ | 14 _Fragment_ | 14 _Fragment_ | 14 _Fragment_ | | 14 _Fragment_ |
| Observations | 57 | 57 | 57 | 57 | 48 | 48 | | 48 | | 48 | 126 | 126 | 126 | | 126 |
| Marginal / Conditional R^2^ | 0.128 / 0.535 | 0.171 / 0.550 | 0.219 / 0.594 | 0.326 / 0.564 | 0.412 / 0.783 | 0.251 / 0.551 | | 0.112 / 0.406 | | 0.547 / NA | 0.731 / 0.949 | 0.689 / 0.961 | 0.643 / 0.926 | | 0.296 / 0.927 |
| AIC | 232.409 | 180.939 | 115.053 | 72.880 | 329.426 | 336.005 | | 271.931 | | 292.561 | -484.485 | -490.838 | -430.210 | | -189.011 |

C. **Figures**

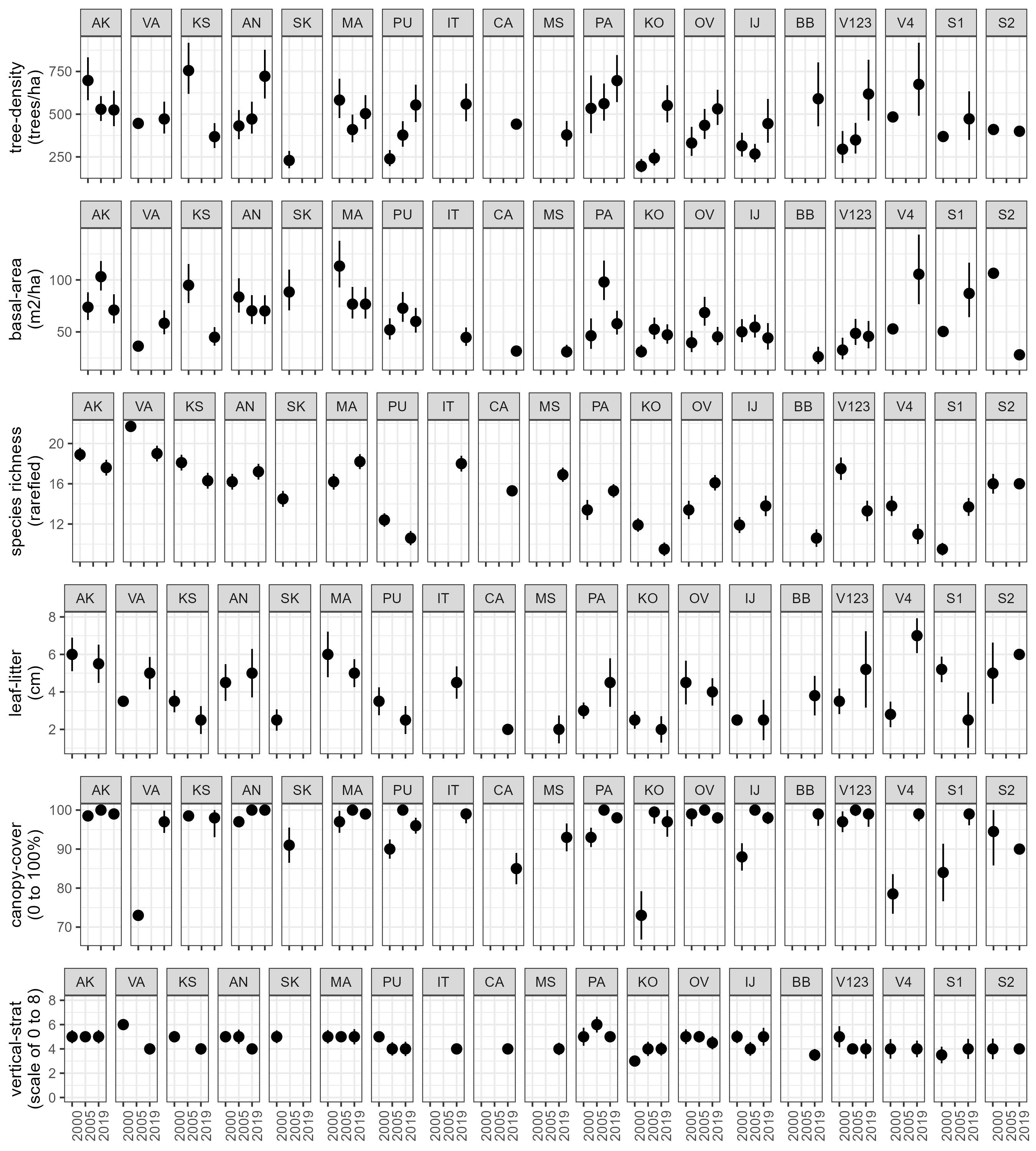

**Figure S1:** Variation in habitat structural parameters across sites and seasons. Site codes as in Table 1.
